# LOESS and DE-SWAN can induce artifactual “waves” of molecular aging

**DOI:** 10.64898/2026.06.24.734079

**Authors:** Madeleine Carbonneau, Katherine H. Shutta, Jeffrey W. Miller, Michael P. Snyder, Xiaotao Shen, John Quackenbush

## Abstract

A growing body of literature has investigated the relationship between age and biomolecular changes, leading to conclusions that aging occurs in discrete molecular “waves.” Data summary tools such as LOESS and sliding window analyses like DE-SWAN are common approaches that have gained acceptance in recent years. We demonstrate via simple simulations that these tools can identify non-linear patterns of aging where they do not exist. Specifically, we show that (i) clustering of molecular trajectories using LOESS can lead to artifactual characteristic patterns of molecular aging, (ii) “waves” of aging identified using the combination of LOESS and DE-SWAN in real data are not robust to changes in the underlying age distribution and are not supported by valid permutation testing, and (iii) DE-SWAN alone can generate pronounced “waves” of nonlinear molecular aging in linear data due to differences in statistical power along the age continuum. Our results specifically challenge the statistical support for discrete aging crests inferred in the literature, but do not rule out nonlinear molecular aging or age-associated transitions that may be detectable using other cohorts and statistical models.

## 1 Introduction

An increasing number of studies have explored aging at a molecular level with the goal of identifying biological processes that change over time. Such studies typically assay multi-omic abundance data (such as gene expression, protein expression, and metabolomic profiles) in cohorts of individuals spanning a broad range of ages. These data are then analyzed to identify molecules that change along the continuum of age, with an emphasis on finding non-linear trajectories. Molecules showing significant differences are further examined to generate hypotheses about how age-dependent alterations may influence health and disease (17; 26).

Aging and its related phenotypes are challenging to study for a variety of methodological reasons that include survivorship bias, unmeasured confounding, high prevalence of multimorbidities, and long latency periods (21). First, it is often impractical to sample uniformly across the age range being studied, and statistical power to detect molecular changes is generally greater in regions of the age continuum with more participants, typically middle age. Second, even when researchers achieve more uniform sampling, survivorship bias remains a concern: older participants represent a selected population of survivors. Finally, aging cohorts often exhibit age-dependent differences in other important data characteristics, including variance and the prevalence of outliers. These differences can further affect statistical power in an age-dependent manner and complicate the interpretation of age-associated molecular changes. Together, these and other factors make it difficult to accurately model the relationship between age and biomolecular expression across the lifespan. Longitudinal cohort-based studies (such as the Baltimore Longitudinal Study of Aging (8), the Dunedin Multidisciplinary Health and Development Study (24), and the Study of Muscle, Mobility, and Aging(4)) partially mitigate some of these issues. However, for both cross-sectional and longitudinal data, differences in statistical power across the age continuum make true molecular behavior difficult to characterize.

Many recent studies of aging use locally estimated scatterplot smoothing (LOESS) (3), Differential Expression - Sliding Window ANalysis (DE-SWAN) (17), or a combination of the two. LOESS (an acronym for LOcally Estimated Scatterplot Smoothing) is a nonparametric regression method used to estimate smooth relationships between variables without assuming a specific functional form. Instead of fitting one global curve (like a straight line or polynomial) to all the data, LOESS fits many small local regressions around each point and assembles them into a smooth curve (3; 12). At each point, the local regression function is estimated using a subset of the data belonging to a local neighborhood. The proportion of the data included in this neighborhood is a tuning parameter called the “span” tuning parameter, which can be optimized via cross-validation. Typically, each data point is weighted so that points closest to the neighborhood center have a larger influence on the local regression. After the local regressions have been fit at each point in a grid, the fitted values are connected to create a smooth curve (12).

The Differential Expression-Sliding Window ANalysis (DE-SWAN) algorithm uses a sliding window approach to detect points at which molecules are differentially expressed along a quantitative trait, which is typically age (17). Within each window, the method tests for a difference between individuals younger and older than the window center (Figure 1). This method has been deployed across many molecular assays (such as RNA-seq, proteomic, and metabolomic data) so that results from the collection of differentially expressed molecules can be pooled to identify particular ages at which there may be a concerted biomolecular change (26).

**Figure 1:**
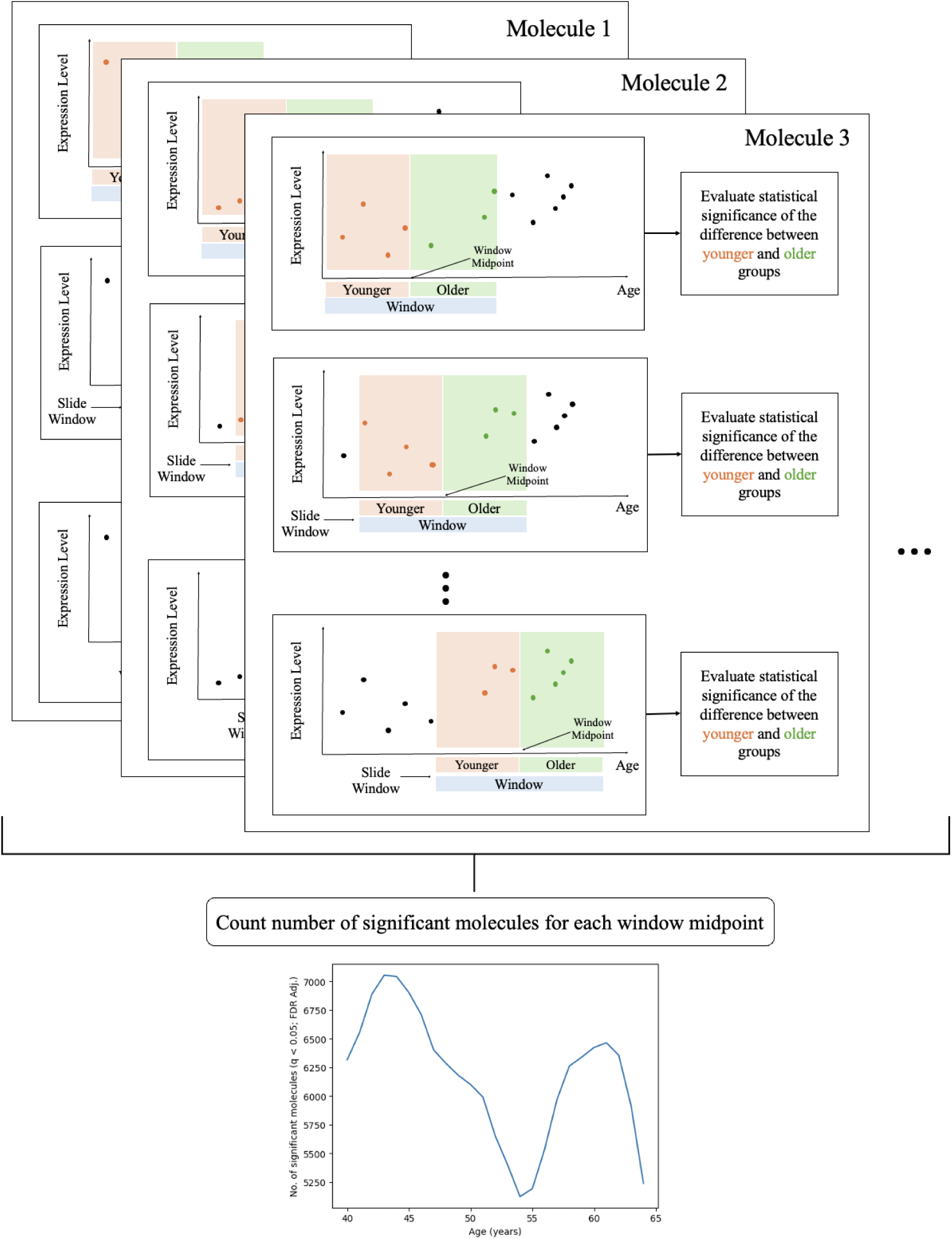
Example of DE-SWAN algorithm using toy data. The graphic shows how, for each molecule, the sliding window analysis traverses the age range. The orange window shows the younger group and the green window shows the older group. At each window midpoint, the difference between these two groups is assessed using a statistical test. The number of molecules assessed as significantly different at each window midpoint is then plotted and interpreted for patterns such as nonlinearity.

DE-SWAN is based on two key input parameters: a set of window centers and a set of “bucket” sizes determining the width of each window. Based on these choices, DE-SWAN sequentially steps through the window centers and applies a statistical test for differential expression between two groups: those samples falling in the lower half of the window (in the case of age, “younger”) and those in the upper half (“older”). Repeating this test for each molecule results in a set of molecules determined to be significantly differentially expressed between the two halves of the given window. The algorithm then shifts to the next window center and repeats the process (Figure 1). The original DE-SWAN algorithm uses a linear regression model to compare expression between the younger and older groups (17). However, DE-SWAN has been adapted to test for differential expression using other statistical tests (26). Importantly, in its conception, the statistical properties of DE-SWAN were not rigorously assessed (17). As a result, there has been no comprehensive evaluation of its behavior in large-sample and finite-sample settings. Further, because DE-SWAN is a method built from per-timepoint measures of significance instead of assessing association along the age continuum as a whole, it can elicit conclusions of nonlinearity that are driven by structural features of the data (such as the age distribution, heteroskedasticity, and outliers) rather than by substantive biological signal (such as true changes in expression).

Here, we critically analyze how LOESS, DE-SWAN, and the combination of these methods have been used to study the aging process. We show that when applied to null datasets consisting of random noise, these tools can identify strong, dramatic patterns that mirror those in published papers analyzing the effects of aging. Further, the outputs from these methods are not actually a measure of association with age; a nonlinear number of significant tests as a function of age does not imply a nonlinear association between age and the abundance of a molecule or classes of molecules. Absent unfeasible assumptions about structural features of the sampled data (such as constant sample density, variation, outlier density, etc. across the age distribution), the results of these pipelines cannot reliably characterize the aging process.

We note that while we focus on LOESS here, LOESS is merely one instance of a broader class of statistical tools for smoothing and interpolating data, such as polynomial regression, spline modeling, or generalized additive models (GAMs). The use of any such tool could potentially lead to artifacts of the type described below. The key methodological error that yields artifactual waves is not LOESS per se, but rather the coupling of a smoothing method with downstream analyses without appropriate propagation of uncertainty.

The remainder of this manuscript is structured as follows. In Section 4.1, we discuss established weaknesses of LOESS and show how these weaknesses can be problematic for the common “smooth-cluster-interpret” workflow used in recent papers (for example, 5; 17; 18; 23; 26; 31; 32). In Section 2.2, we show how the combination of LOESS and DE-SWAN can find artifactual “waves” of aging in simulated datasets that were explicitly constructed to have no true signal. We also discuss how LOESS+DE-SWAN has been used to find waves in real datasets and how these identified waves do not survive permutation testing (26). In Section 2.3, we show that DE-SWAN (without LOESS) can indicate evidence of nonlinearity even when the underlying data-generating process is strictly linear, due to variations in statistical power at different ages. Lastly, in Section 3, we discuss the importance of these findings and provide recommendations for future analyses of molecular aging and other related life course data. In Section 4.4, we provide a brief assessment of the prevalence of these issues in the aging literature. All code is open-source and publicly available at https://github.com/QuackenbushLab/artifactual-waves-of-aging.

## 2 Results

### 2.1 Artifact 1: LOESS can induce clustered trajectories in null data

It has become common in the aging literature to model continuous trajectories of molecular levels as a function of age by generating smooth approximations (often using LOESS) from discrete measurements of these molecules originally taken from a cross-sectional population. In a typical workflow, these smoothed trajectories are then clustered, and cluster centroids are interpreted as characteristic profiles of aging (for example, molecules that are stable through midlife and then increase in expression or molecules that are highly expressed in younger individuals and then decrease with age). Enrichment analyses are typically conducted on these clusters to identify biological functions that may exhibit these characteristic trajectories through the life course. We refer to this as the “smooth-cluster-interpret” workflow. We conducted simulations that demonstrate that the smooth-cluster-interpret workflow can find “biologically interpretable” aging trajectory shapes in null data.

We simulated uniform, Gaussian, and Gaussian mixture age distributions (*n* = 200 samples in each case) as well as the age distribution underlying the transcriptomic results of Shen et al. (26) (*n* = 97); these data are part of the Integrated Personal Omics Profiling (iPOP) project Chen et al. (2) and are hereafter referred to as the “iPOP age distribution.” For each sample, we simulated null expression levels for 1000 molecules, where “null” refers to data generated from a standard Gaussian distribution that is independent of age. We then applied LOESS to these data and clustered the resulting trajectories (Online Methods 4.1).

The age distributions, raw data, and LOESS-smoothed data are shown in Figure 2A. As expected, no apparent trajectories are observed in the raw data, and clustering yields a dendrogram of nearly equal height for each predictor. After applying LOESS to the data, smooth trajectories appear in the heatmap and a clear clustering structure appears in the dendrogram. The case of the uniform distribution serves as a control in this study: centroids of clusters estimated from the uniform distribution are relatively flat, except for small variations induced by sampling variability (Figure 2B). In contrast, centroids of clusters learned from the Gaussian, Gaussian mixture, and iPOP age distributions exhibit similar trends to those that have previously been presented as true signals in the aging literature (Figure 2B). Because sampling is typically most sparse in the tails of the distribution, there are often fewer points to “anchor” the LOESS curve. We found that LOESS curves tended to deviate from the true flat trajectory at the ends of the age distribution, creating the false impression that molecular expression changes in youth and/or older age but remains stable through midlife. Similarly, for bimodal data, we observed peaks in the cluster centroids around middle age, suggesting molecular changes specific to midlife that were not actually present in the underlying data. Furthermore, as the number of clusters increases, the signals often become more “interpretable” – a conclusion that is, in fact, due to overfitting (Supplementary Figure **??**).

**Figure 2:**
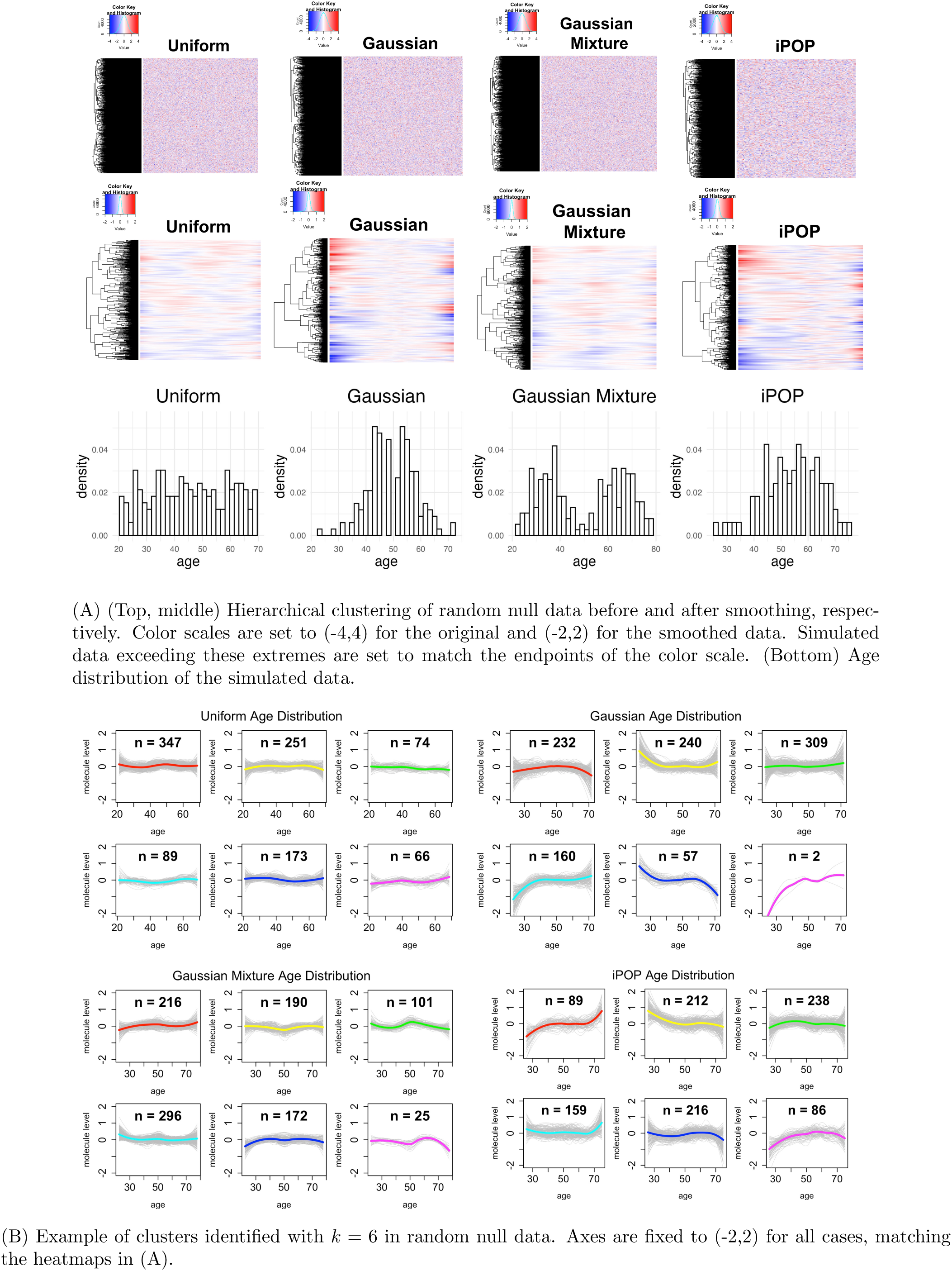
Hierarchical clustering of random null data before and after smoothing shows that smoothing induces patterns of aging that follow the age density, even when there is no true signal.

In our analyses, we used LOESS for smoothing because it is the method most commonly used in published studies of molecular aging. However, the artifacts we observed are not necessarily unique to LOESS and could also arise with other smoothing methods, including moving averages, splines, generalized additive models (GAMs), and Gaussian filters. The “smooth-cluster-interpret” workflow is particularly problematic when applied to the sparse, non-uniform age distributions common in molecular aging studies. Our results demonstrate the importance of including appropriate null comparisons when using this workflow. Without such comparisons, it is difficult to determine whether an observed pattern reflects a genuine biological signal or is simply an artifact of the analysis.

### 2.2 Artifact 2: LOESS+DE-SWAN can elicit artifactual waves of aging that are not supported by null simulations and permutation testing

The LOESS+DE-SWAN workflow combines LOESS with DE-SWAN by first applying LOESS to the original data to interpolate a smooth curve of molecular expression and then running DE-SWAN on these interpolated values, using a Wilcoxon test to compare younger and older groups in each window (Figure 3A) (26). Using LOESS and DE-SWAN simultaneously is problematic because the points on a fitted LOESS curve are not independent from one another. Instead, each point along the curve is estimated relative to the same dataset and constrained to follow a smooth trajectory. Therefore, points along this curve will be highly dependent and cannot be used as independent observations supporting the same trend. Using LOESS-interpolated data with DE-SWAN thus yields an immediate concern: a statistical test assuming independent data points is instead being used on highly correlated data. Most tests assume independent data points, so the problem with LOESS+DE-SWAN is not solely due to the use of the Wilcoxon test. However, it is exacerbated by the Wilcoxon test’s rank-based nature; the smoothness of the LOESS-interpolated curve makes the rank ordering often nearly monotonic from the left to the right side of the sample window. We illustrate this phenomenon in Section 2.2.1 below.

**Figure 3:**
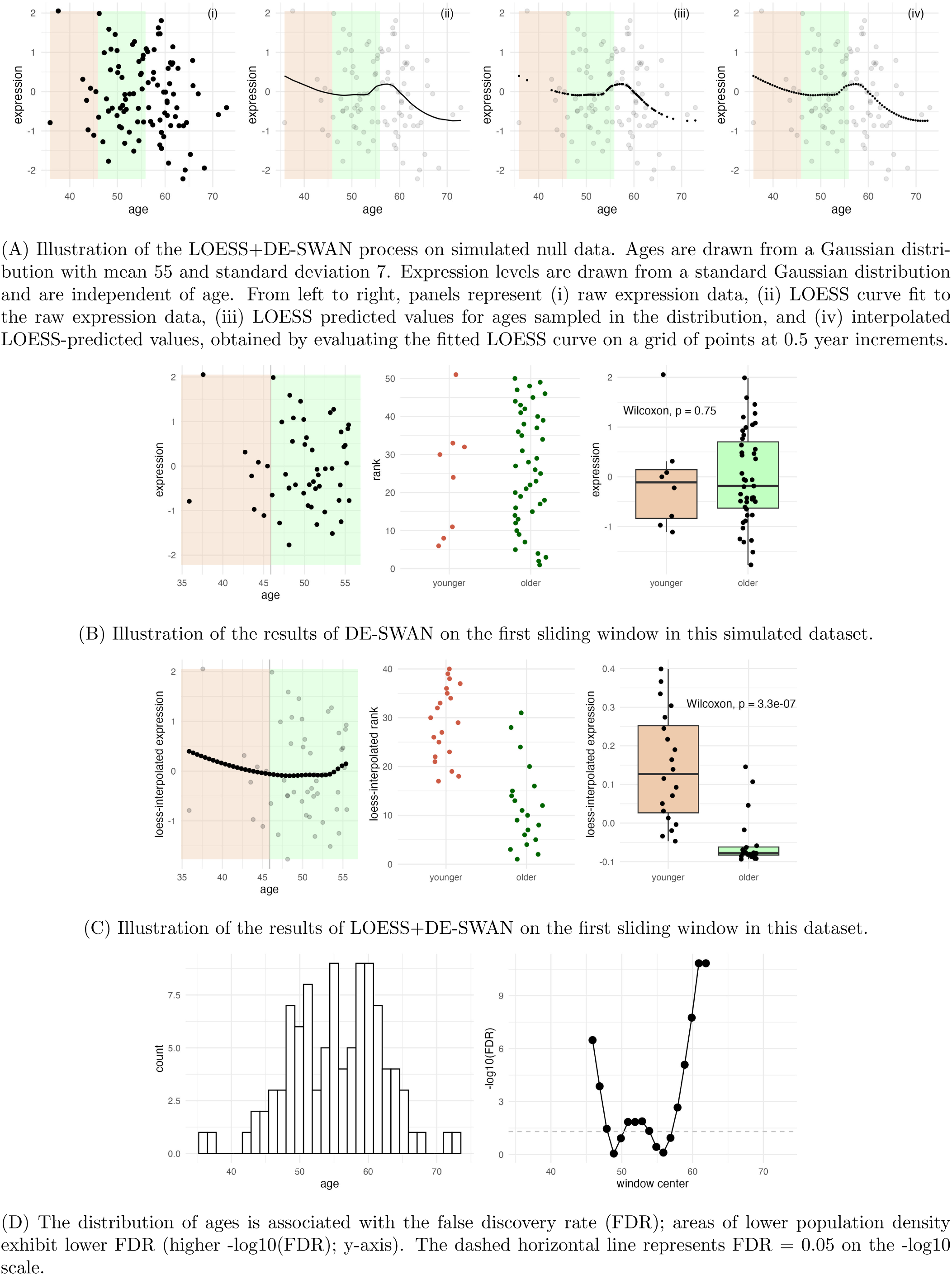
A simulated example of inflated detection of differential expression in LOESS+DE-SWAN.

#### 2.2.1 Simulation example on a single gene

To demonstrate inflation of significance by LOESS+DE-SWAN, we considered a simple example in which LOESS+DE-SWAN is applied to a single gene with expression levels that have no relationship with age (linear or non-linear) (Online Methods 4.1.1).

As expected, a scatterplot of expression versus age showed no clear relationship (Figure 3a, i). The LOESS curve fit to these data reflects a classic limitation of LOESS: the curve is sensitive to the density of the data distribution, and its tail behavior differs from the behavior at the center due to the lower data density at the margins (Figure 3a, ii). When the LOESS curve was used to estimate expression according to the original age distribution, an artifactual trend was observed in the random data (Figure 3a, iii). This artifactual trend was amplified when LOESS was used to interpolate data points across a uniform grid of points positioned every 0.5 years on the age distribution (Figure 3a, iv).

While the scatterplot visualization already demonstrates how LOESS+DE-SWAN identifies false signals, direct statistical calculations reveal the magnitude of this issue. Figure 3B shows an example of DE-SWAN without LOESS. In this case, the ranks of the expression data are similarly distributed between the “younger” and “older” groups, as reflected by a non-significant Wilcoxon test result (p=0.75). Figure 3c shows the same calculation performed on the LOESS-interpolated expression values. Because of the tail behavior of the LOESS curve, the ranks are now substantially higher in the younger group than the older group, and the Wilcoxon test indicates high significance of this difference (p = 3.3 *×* 10*^−^*^7^).

While we focus on the first window in this example, the inflated statistical significance of LOESS+DESWAN appears to be sensitive to the underlying age distribution and persists across many windows in the sliding window analysis (Figure 3D). Windows including data near sparser regions of the age distribution seem to be particularly prone to finding spurious signals in null data.

#### 2.2.2 Waves of aging appear in null data and depend on the age distribution

To illustrate how LOESS+DE-SWAN can generate spurious results related to the underlying age distribution, we conducted a simple simulation study in which we drew expression levels for 1000 molecules, independently of age, and ran LOESS+DE-SWAN to identify “waves” of aging (Online Methods 4.2.1).

Across all four age distributions considered, LOESS+DE-SWAN detected a majority of the 1000 simulated expression profiles to be significantly changing (controlling FDR at 0.05) at each window center (Figure 4). Because there is no true change in the mean expression, these targets represent hundreds of false discoveries per time point. While this is notable in itself, the primary concern with these results is the observed relationship between the number of significant molecules and the age distribution (Figure 4A-B). In particular, when there are few points in one or both halves of the sliding window, the influence of LOESS-interpolated points is amplified. In contrast, DE-SWAN without LOESS controls the FDR, finding few age-associated molecules in this setting where simulated expression is unassociated with age (Figure 4C).

**Figure 4:**
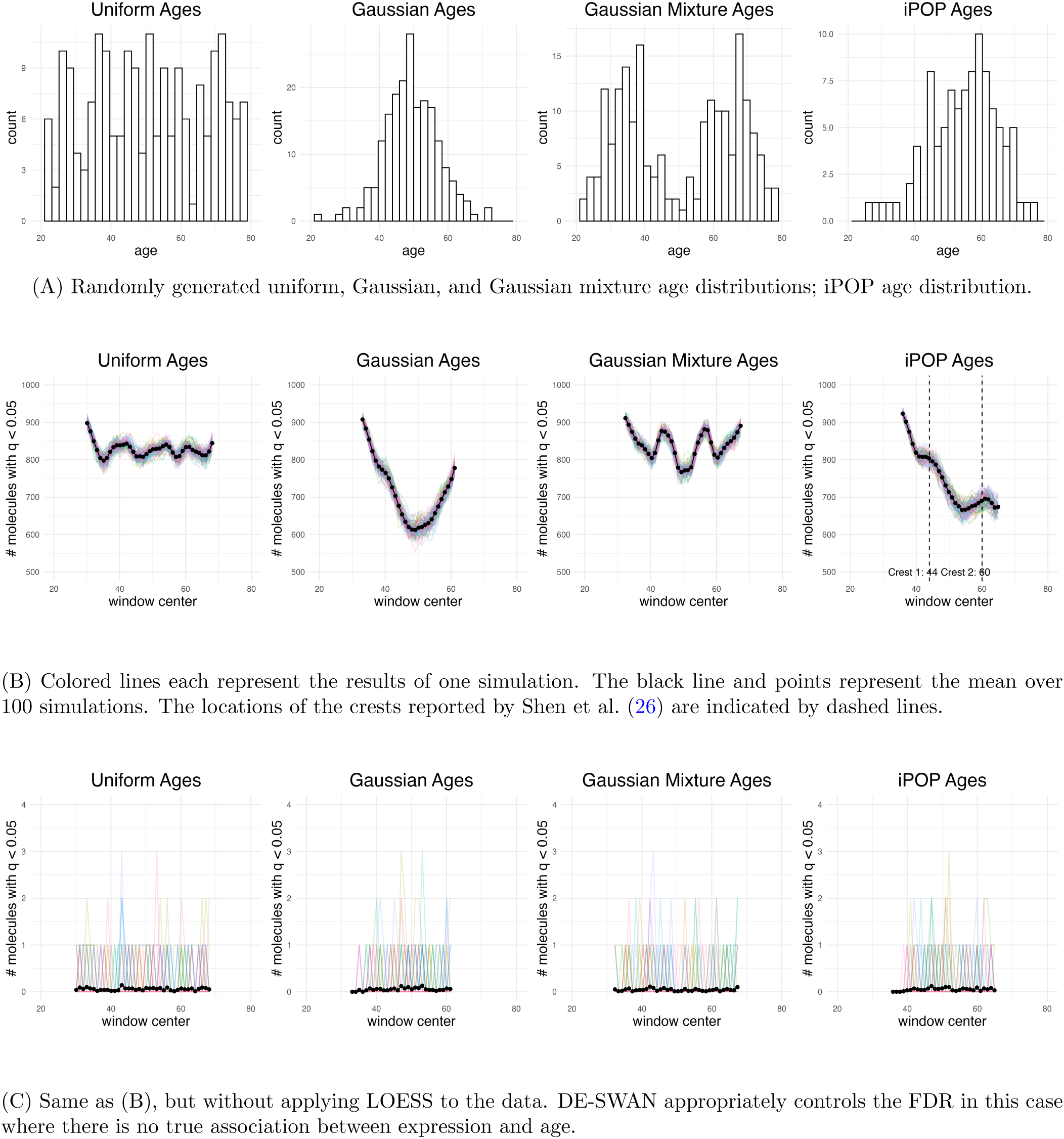
Simulation study of LOESS+DE-SWAN and DE-SWAN on randomly generated data with no true association between molecular expression and age.

We posit that this phenomenon could be the cause of the waves of aging in the mid-40s and early 60s reported by Shen et al. (26). To address this hypothesis more specifically, we conducted an additional simulation that used the iPOP age distribution but simulated random noise as the expression values for each gene, with no true change in mean expression as a function of age (Supplementary Section 4.2.1). Our simulation used the LOESS interpolation procedure (evaluating the LOESS curve at a grid ranging from 26 to 75 with half-year steps) and the same DE-SWAN windows as Shen et al. (26); the initial window is from 25 to 50 with a center at 40, and the remaining windows are of width 20 and centered at 41, 42*,…,* 65.

Figure 5A shows the results of this analysis, with the locations of the transcriptomic crests observed by Shen et al. (26) indicated. While the null results do not perfectly recapitulate the reported iPOP transcriptomic waves, it is notable that despite the lack of any true signal, two crests clearly appear in this null study, one in the early 40s and one in the early 60s. The appearance of two crests at these time points, based on data that is, by design, purely random noise, calls into question the validity of the conclusion that these peaks represent true biological trends—rather than a construct arising from the structure of the population age distribution and the shortcomings of the analytical methods being used. Also notable is the correlation between the sample size in the sliding window and the number of changing molecules that are reported by LOESS + DE-SWAN (Figure 5B).

**Figure 5:**
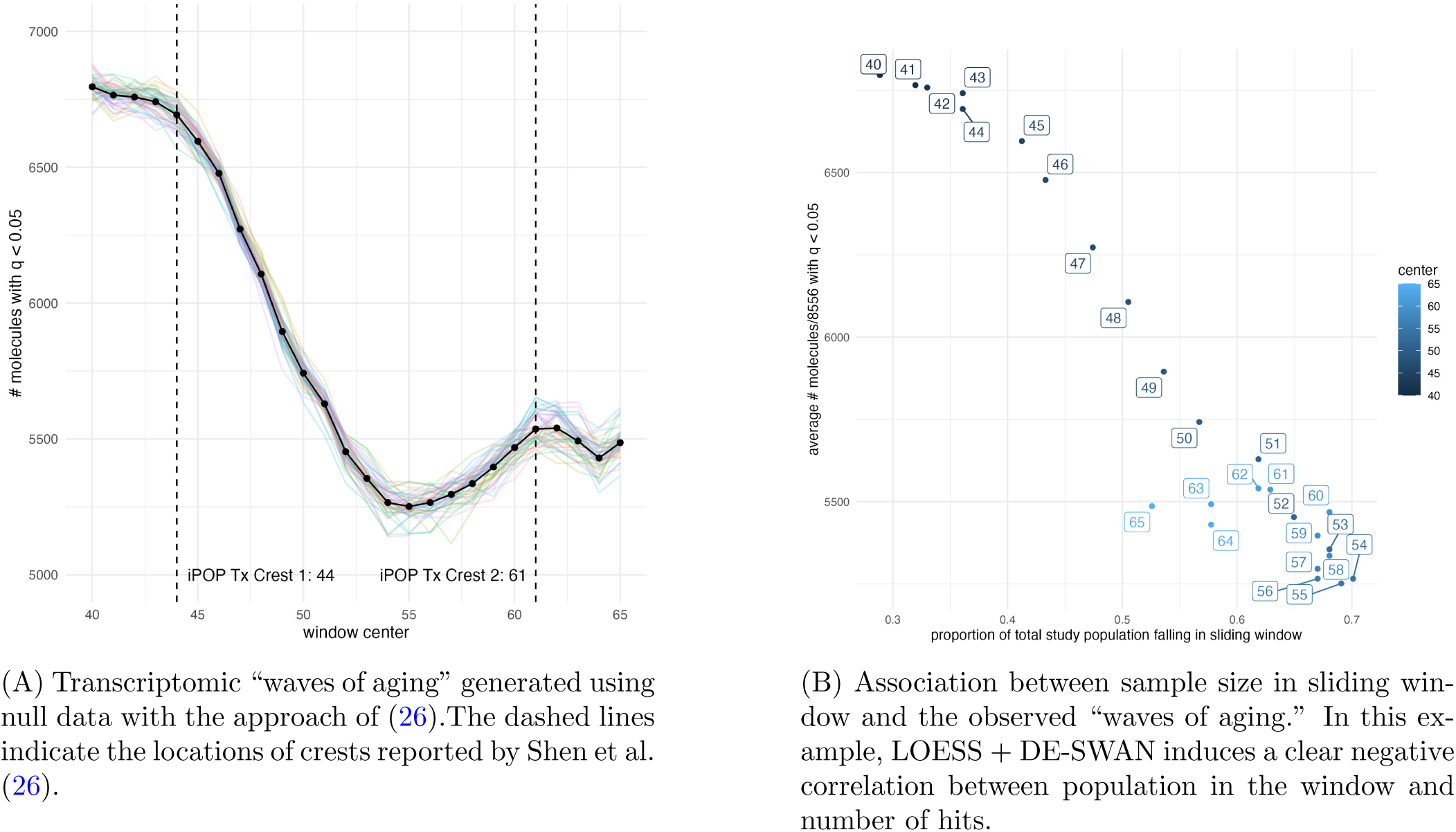
Simulation study repeating the transcriptomic analysis of Shen et al. (26) using the iPOP age distribution but with simulated gene expression values having no true association with age.

#### 2.2.3 DE-SWAN alone does not detect “waves of aging” in the iPOP data

To further examine the role of LOESS interpolation in these results, we analyzed the transcriptomic data from iPOP using DE-SWAN without LOESS. We downloaded code and data published by Shen et al. (26) and reproduced the results shown in Figure 4(d) of Shen et al. (26); see Figure 6A. To investigate the impact of running LOESS before conducting the DE-SWAN analysis, we next ran DE-SWAN on the transcriptomic data without applying the LOESS interpolation. LOESS grid interpolation values were identical to those of Shen et al. (26); DE-SWAN windows were set to width 20 ranged from 40 to 65 in 1-year increments, which closely follows the approach of Shen et al. (26). With DE-SWAN alone, the peaks seen in Figure 6A disappeared; no transcripts met significance after controlling the FDR at 0.05 (Figure 6B).

**Figure 6:**
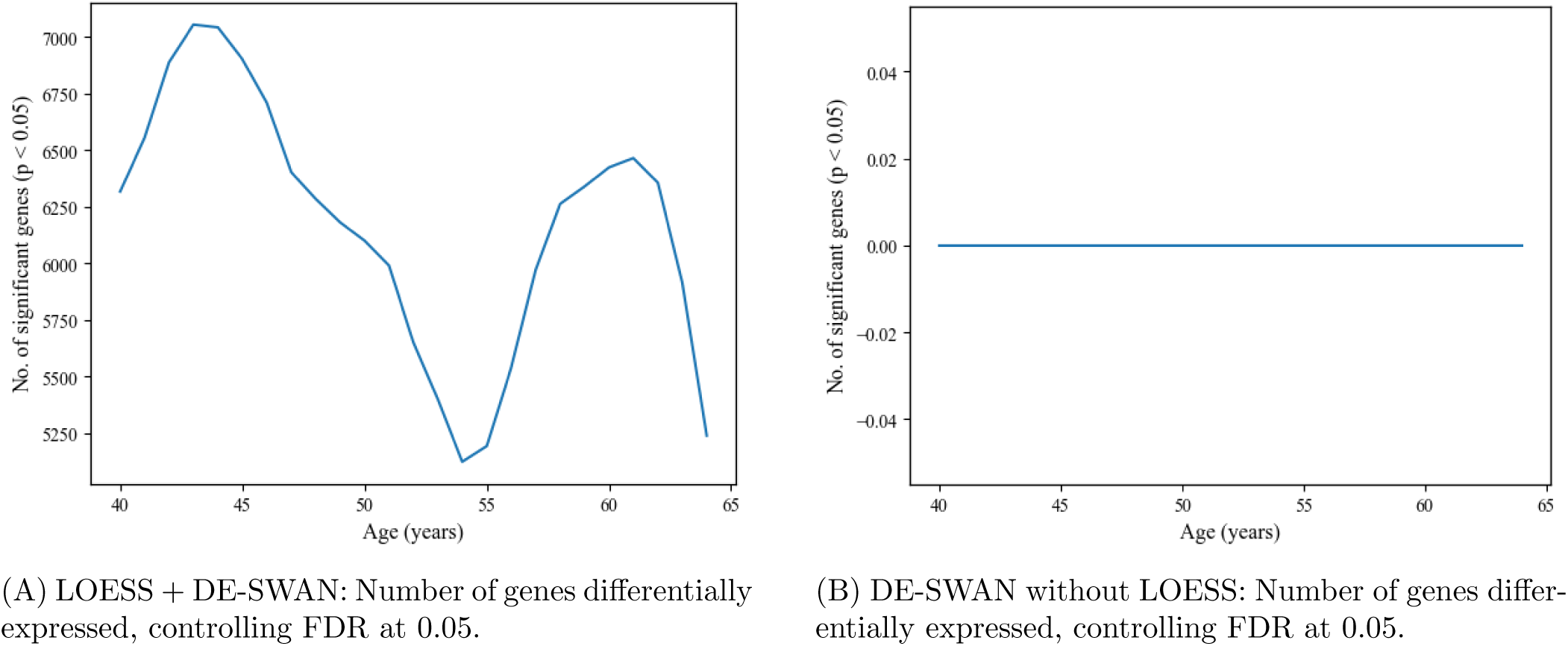
Reproducing the transcriptomic analyses of Shen et al. (26) on the original iPOP data using LOESS+DE-SWAN, and comparing with results obtained using DE-SWAN alone.

#### 2.2.4 Use of permutation testing to validate the procedure

Shen et al. (26) reported the results of a permutation test intended to validate the robustness of the aging crests by preserving the marginal distribution of the expression data while breaking the dependence on age. Based on this permutation test, (26) found that their observed peaks disappeared following permutation of the ages, interpreted as evidence of validity. We attempted to reproduce this result by conducting a permutation test using the method in Algorithm 1 on the iPOP expression data and ages. In contrast to the results of Shen et al. (26), we found that clear peaks remain visible with the permuted data (Figure 7A). Thus, even when we broke the dependence between age and expression in the original data by permuting the ages, LOESS+DE-SWAN still spuriously detected significant peaks that resembled the results from the unpermuted data. These results provide additional evidence that, under the LOESS+DE-SWAN workflow, the peaks observed in Figure 6A cannot be reliably distinguished from artifacts generated by the interaction of smoothing, interpolation, random variation, and the underlying age distribution.

**Figure 7:**
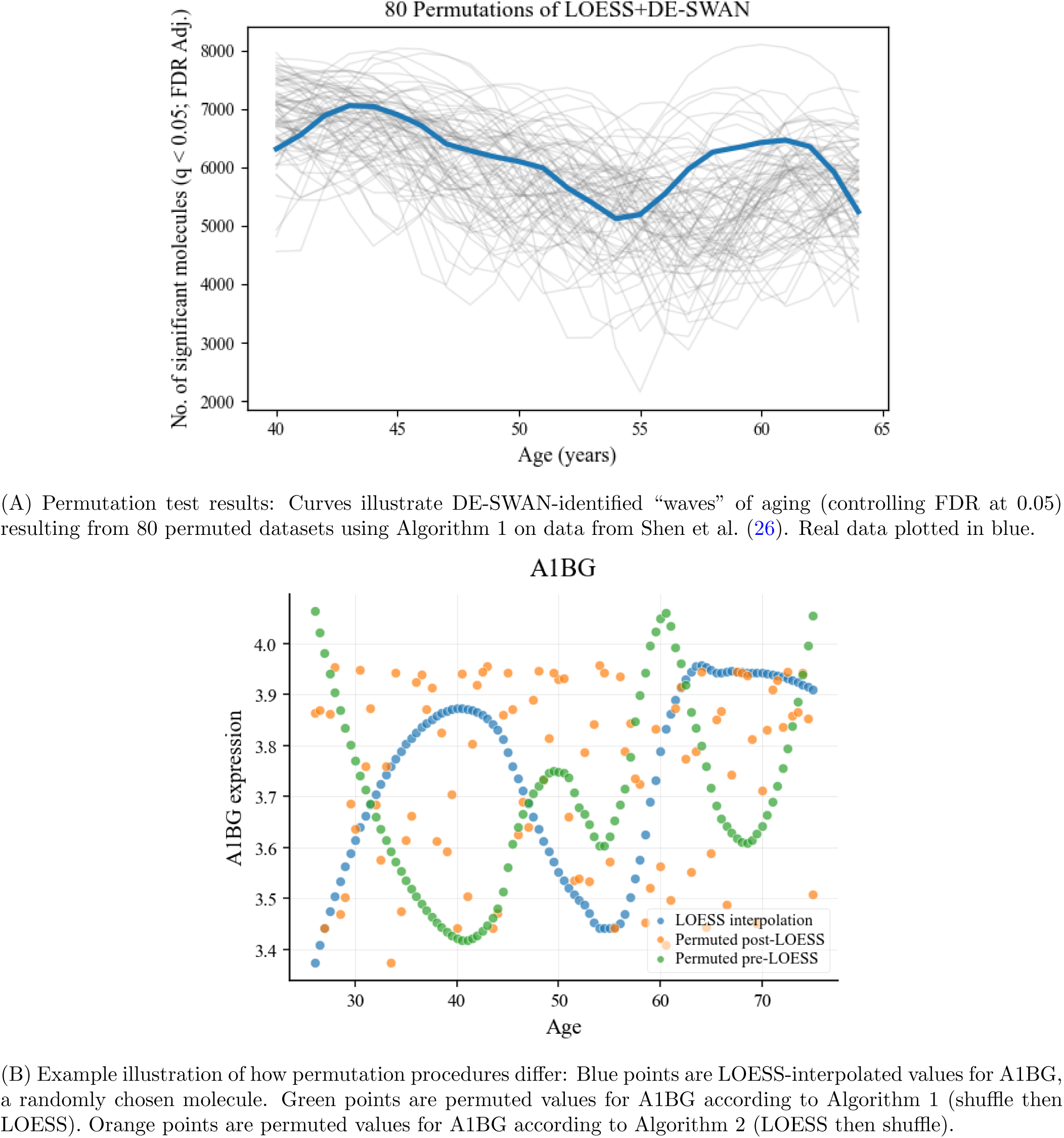
Permutation results and analysis.

##### Algorithm 1 Permutation testing for LOESS+DE-SWAN: Shuffle **before** LOESS

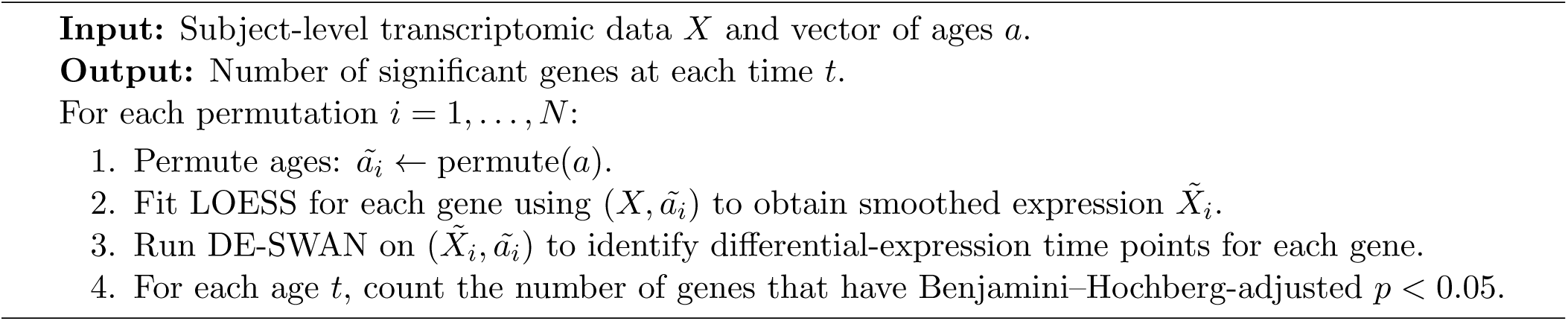

The discrepancy between our permutation results (Figure 7A) and the permutation results reported in Supplementary Figure 4(c) of Shen et al. (26) is that the latter employed a subtly—but importantly— different approach (Algorithm 2). The key difference is that Shen et al. (26) permuted the ages after LOESS interpolation of the data, whereas we permuted the ages before LOESS. This seemingly small difference in procedure produces an invalid null comparison for the question at hand.

##### Algorithm 2 Permutation testing for LOESS+DE-SWAN: Shuffle **after** LOESS

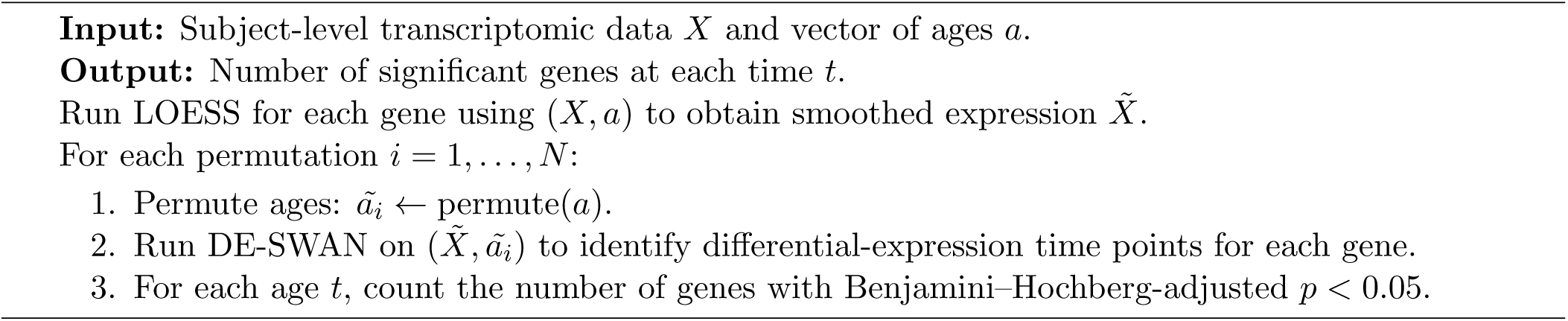

The difference between Algorithm 1 and Algorithm 2 is shown for one gene, A1BG, in Figure 7B. In this figure, we see that the green points (permuted data according to Algorithm 1) are much more likely to elicit positive results from DE-SWAN than the orange points (permuted data according to Algorithm 2).

The validity of permutation tests emerges from the assumption of exchangeability (9). After applying LOESS to the data, points along the LOESS curve are not exchangeable with respect to age. Meeting the exchangeability assumption requires shuffling of the transcriptomic matrix *X*, not the LOESS-interpolated values *X̃*. Algorithm 1 is therefore the appropriate permutation-based approach for assessing the validity of the results shown in Figure 6A. We conclude that multi-omic “waves of aging” presented in Figure 4(d) of Shen et al. (26) are not adequately supported by the original permutation analysis and rather are strongly influenced by noise captured and amplified by LOESS+DE-SWAN.

### 2.3 Artifact 3: DE-SWAN identifies non-linear patterns of aging in linear data due to age-associated differences in statistical power

DE-SWAN identifies age-specific molecular changes by counting the number of significantly changing molecules at each of a sequence of time points along the age continuum (Figure 1). This count of molecules at each age point is influenced by both the true number of molecules whose expression is changing and the statistical power to detect them from the observed data. It is well-established that power is influenced by several characteristics of the data: the sample size, the significance level (*α*), the noisiness of the data, and the magnitude of the true effect (25). To interpret a DE-SWAN curve as a faithful reflection of age-related molecular changes, we must assume that statistical power depends only on the magnitude of the true effect, which is unlikely if the other factors vary with age. Such an interpretation must also be qualified by the fact that the DE-SWAN curve represents only the number of significant molecules per timepoint, and does not represent the actual association of a particular molecule or set of molecules with age.

Here, we illustrate the potential of differential statistical power to induce artifactual waves of aging by conducting a simulation study in which molecular expression changes linearly with age, so the magnitude of the true effect is constant. We show that non-uniform age distributions (Section 2.3.1), age-dependent heteroskedasticity (Section 2.3.2), and outliers (Supplementary Section **??**) can produce DE-SWAN results that indicate non-linear patterns, even though the true change is linear.

In the present simulation study, we simulated the true molecular expression as linearly changing with age. Therefore, any nonlinearity shown in DE-SWAN curves does not result from true nonlinear expression, but rather some other aspect of the data. Specifically, we simulated the expression *Y_ij_* for gene *j* in subject *i* by sampling from the model:

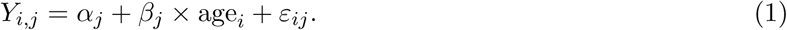

Mechanisms for specifying *α_j_*, *β_j_*, age*_i_*, and *ε_ij_* vary among simulations (Online Methods 4.3).

#### 2.3.1 Sample age distribution

When the DE-SWAN algorithm is applied to a non-uniform age distribution, some age ranges will have more observations than others as the sliding window traverses the age distribution. This change in the number of samples per window results in variation in the power to detect changing molecules at each age. This is illustrated in Figure 8, which shows that different age distributions generate profoundly different DE-SWAN results on simulated linear data. For each distribution and simulation, the data were simulated according to the same procedure (see 4.3). Therefore, the differences in these DE-SWAN results are attributable solely to the differences in age distribution.

**Figure 8:**
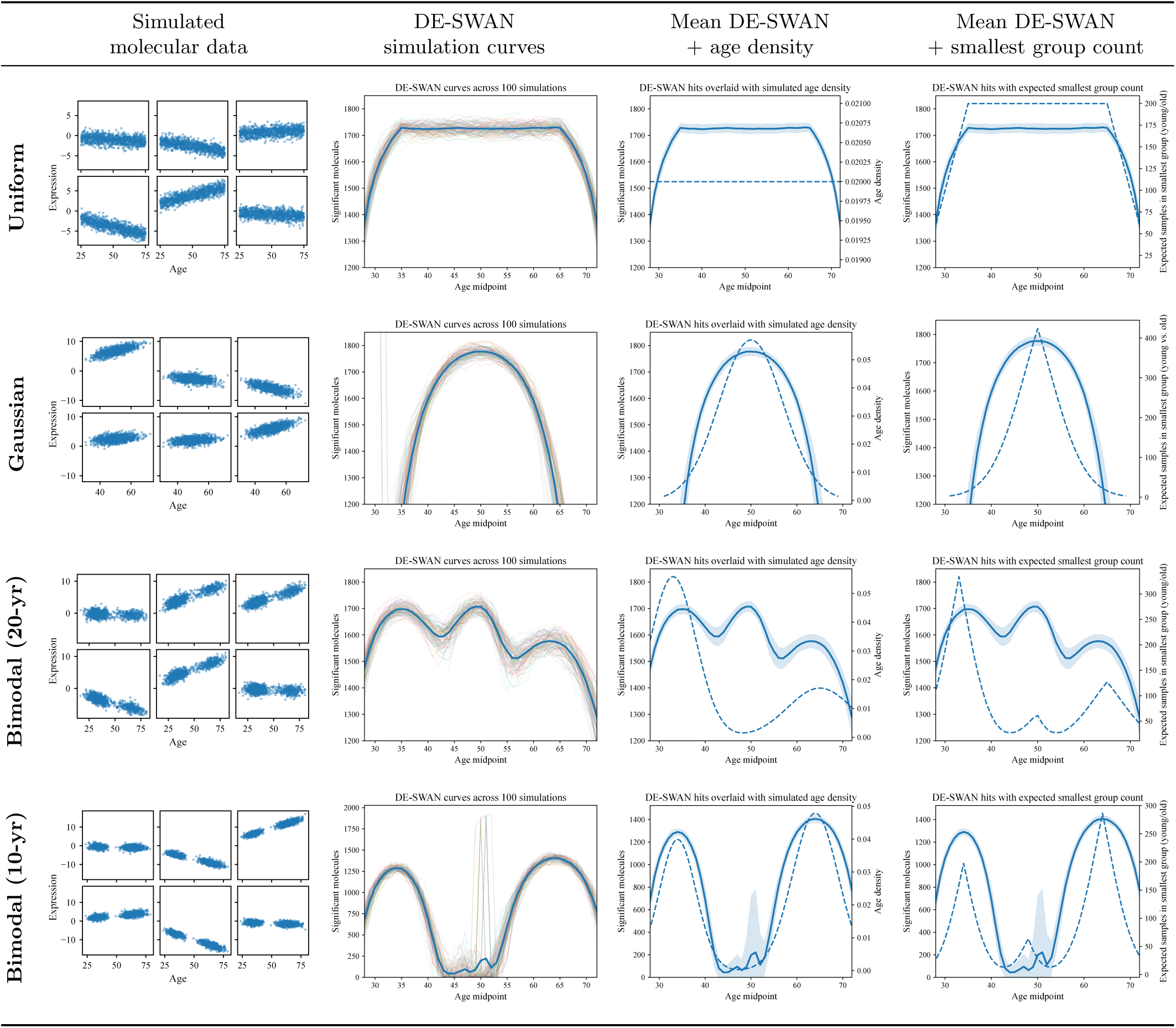
Effects of modulating age distribution on DE-SWAN results. Each dataset was simulated for 1000 observations and 2000 molecules. Results from 100 simulations are shown. The first column shows six randomly drawn molecular expression profiles from a single simulation for each of the population age distributions. The second column shows DE-SWAN curves plotted for 100 simulations for uniform, Gaussian, and two specifications of a bimodal (Gaussian) distribution. The mean of the curves is plotted in blue. The third column shows the mean DE-SWAN curve plotted with a *±*1 SD band (solid blue). The population age distributions (from which each simulation’s age distribution is drawn) are plotted in the blue dashed line. The last column shows the same mean DE-SWAN curve plotted against the expected number of samples per group (older vs. younger) for the group with the smallest number of samples for each slide of the window. Simulation details can be found in Online Methods 4.3.

One tempting “solution” to this problem is to use quantile-based window centers to try to minimize variation in age group size; indeed, this is the default setting for the window parameter in the DE-SWAN R package (16). However, this does not ensure that all statistical tests made along the age distribution have the same number of samples in each group. Because a constant window length is still used, the endpoints of the window do not necessarily coincide with the selected quantiles. The choice of the window length and the midpoint locations dictate which samples will be included in each statistical comparison between younger and older groups, and quantile-based window centers alone will not guarantee that each group contains the same number of subjects.

Even if techniques such as quantile-based window centers, variable window lengths, or resampling methods are used to ensure that each window has the same number of samples, additional mechanisms related to the age distribution can impact power.

We also observe that the intra-window sample density impacts statistical power (see Figure 9). For all three panels, the linear relationship between age and expression is the same. However, when samples are closer to the center of the window (middle panel), the difference between the younger and older groups is smaller than if the samples are located near the boundaries of the window (last panel). In this way, the distribution of the samples within the sliding window impacts the effect size of the difference between the two groups. When samples are located near the boundaries of the window, there will be higher statistical power to detect this change (*p* = 2.3 *×* 10*^−^*^56^) than when samples are located at the center of the distribution (*p* = 1.3 *×* 10*^−^*^17^). This may also help explain the middle inflection point for the bimodal (20-yr) distribution’s DE-SWAN curves in Figure 8. At the location of the middle peak, the samples are densest at the boundaries of the window, inducing a larger effect size.

**Figure 9:**
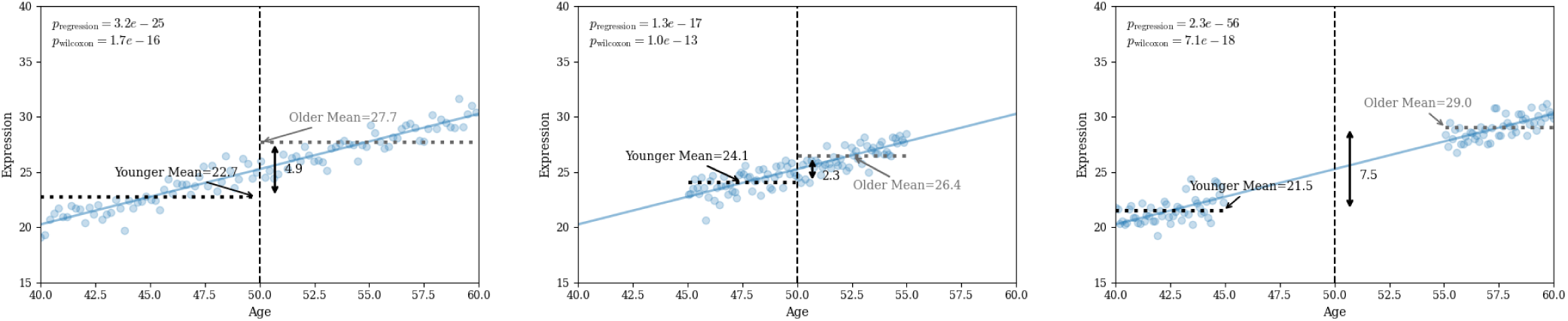
Effects of intra-window age distribution on DE-SWAN results for one window centered at 50. All three panels show data generated for one simulated molecule: *Y_i_* = 0.5 + 0.25 *×* age*_i_* + *ε_i_* where 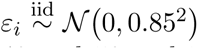. For each panel, there are 100 observations with 50 samples falling between the ages of 40 and 50 and 50 samples falling between the ages of 50 and 60. In the left panel, samples are evenly distributed throughout the window. In the middle panel, samples are concentrated at the center of the window. In the right panel, samples are concentrated at the boundaries of the window. The vertical black dashed line shows the window midpoint (50). The horizontal dotted black and gray lines show the means of the younger and older groups (respectively). The black line with arrows shows the difference between these two means. Two different statistical tests were used to evaluate the evidence of differential expression between the younger and older groups: the regression-based test (*Y_i_* = *α* + *β ×* `1`(Age*_i_ ≥* 50)) (17) or the Wilcoxon rank-based test (26). *P* -values for both of these tests are shown in the top left of each panel.

This helps explain the appeal of using LOESS-interpolated data as input to DE-SWAN. When evaluated on a fixed age grid, the interpolated data appear to be uniformly distributed across the age range (26). However, smoothing cannot change the distribution of the underlying samples or the information they contain. Thus, although combining LOESS with DE-SWAN appears to address the problem of the non-uniformity of the age distribution, it introduces new sources of differential power through other mechanisms (see Section 2.2).

#### 2.3.2 Heteroskedasticity

Heteroskedasticity refers to the presence of non-constant variance along the range of a predictor—in this case, age—and can be problematic for many statistical analyses (7; 10). It is biologically feasible that age can drive heteroskedasticity in molecular expression; in fact, the existence of this phenomenon has been reported previously (14). The presence of non-constant variance in molecular expression data has important implications for DE-SWAN. As variance changes along the age distribution, so does the statistical power to detect molecular change, yielding nonlinearity in the DE-SWAN curves.

We simulated various types of heteroskedasticity in linearly changing molecules to assess the effect of heteroskedasticity on DE-SWAN (Figure 10). In each case, ages were sampled from a uniform distribution and data are generated following the linear model in Equation 1, with the variance of *ε_ij_* depending on age*_i_* (Online Methods 4.3.2). For some molecules, expression might be more tightly or less tightly regulated at the center of the age distribution (Figure 10, first and second rows). For others, expression in older ages might be more variable (Figure 10; third row). Alternatively, some molecules might exhibit more variable expression in younger ages (Figure 10; fourth row). These different trends in variance along the age distribution resulted in different DE-SWAN curve shapes, despite the uniform nature of the age distribution in this simulation. Moreover, it is likely that different molecules will exhibit different variability trends. The overall shape of the DE-SWAN curves will be impacted by the relative contributions of molecules exhibiting a range of patterns such as the ones in Figure 10.

**Figure 10:**
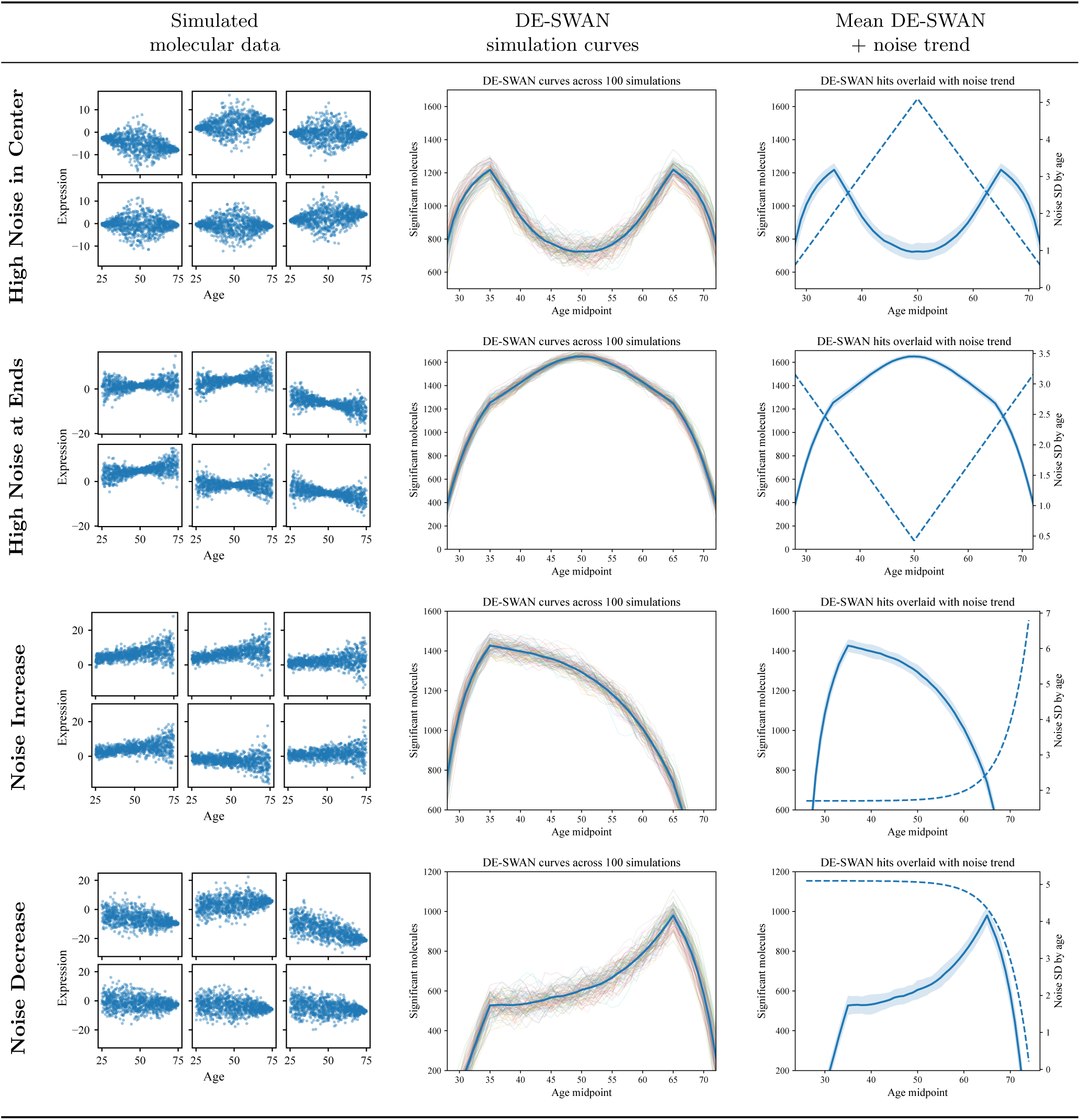
Effects of non-constant noise on DE-SWAN results. Each dataset was simulated for 1000 observations and 2000 molecules, with a uniform age distribution. Results from 100 simulations are shown. The first column shows six randomly drawn molecular expression profiles for one simulation for each variance trend. The second column shows DE-SWAN curves plotted for 100 simulations, with the mean curve plotted in blue. The third column shows the mean DE-SWAN curve plotted with a *±*1 SD band (solid blue). The population standard deviations are plotted as blue dashed lines. Simulation details can be found in Online Methods 4.3.2.

## 3 Discussion

In this manuscript, we presented critical flaws of three techniques that have been applied in many high-impact aging studies: LOESS, DE-SWAN, and the combination of the two (LOESS+DE-SWAN). Our analyses call into question published conclusions about “waves of aging” and non-linearity of the aging process that are based on these tools.

We observed that clustering the output of LOESS on simulated null data produced clusters of molecules with trajectories similar to those reported in the aging literature. Such trajectories have been interpreted as indicating biologically meaningful properties; however, our analyses caution against such interpretations since similar trajectories can appear when there is no true association with age.

Our work shows that DE-SWAN can produce a range of non-linear curves on simulated null data that actually have a linear relationship with age, due to variation in power across age windows.The shape of these curves is strongly affected by non-uniform age distributions, heteroskedasticity, and clusters of outliers, all of which occur in multi-omic datasets. Without accounting for varying statistical power versus age, the shape of a DE-SWAN curve is difficult to interpret. Our simulation studies clearly demonstrate that differences in power can induce artifactual waves of aging in DE-SWAN curves when there is no basis for these patterns in the data. Moreover, there is a risk for misinterpretation of results built using DE-SWAN, arising from a fundamental misunderstanding of what the method measures. DE-SWAN summarizes the results of multiple statistical tests performed along the age continuum, rather than directly measuring the strength or magnitude of associations with age.

Finally, the LOESS+DE-SWAN pipeline can indicate seemingly strong nonlinear signatures of molecular expression. However, when we rigorously tested these findings using null simulations and permutation tests, we did not find any evidence attributing these signatures to true biological change. Instead, we find evidence that applying DE-SWAN to LOESS-smoothed data artificially boosts significance, and that the patterns that emerge appear to be driven by stochastic properties of the data, small numbers of samples, and the underlying age distribution.

This manuscript has been partially motivated by the rapid proliferation of papers that use LOESS, DE-SWAN, or related analytical methods to identify age-related patterns in molecular profiles (see, for example, Ding et al. (5); Eduthan et al. (6); Lai et al. (15); Lehallier et al. (17); Liu et al. (18, 19); Shen et al. (26, 27); Yin et al. (31); You et al. (32)). As of September 26, 2026, Google Scholar reports more than 1100 citations of (17), which introduced DE-SWAN and popularized the “smooth-cluster-interpret” workflow. Integrating query results from three databases (Google Scholar, Scopus, and ScienceDirect) and manually screening the results, we identified 63 papers, 18 of which were published in 2025 and 25 of which were published 2026, that (1) cite (17), (2) use LOESS and/or DE-SWAN in their methodology, and (3) focus on aging or age-related diseases as a primary topic (Supplementary Section 4.4, Supplementary Figure **??**). In many cases, it was not possible to confidently answer (2) in the affirmative or negative. The prevalence of this uncertainty is a key part of the issue.

Disentangling signal from artifact in conclusions based on statistical methods that do not adequately discriminate between differential statistical power and true molecular change is a difficult task. Statistical power is an elusive quantity that can be influenced by many aspects of the data and the methods that are used in its analysis. Without a reliable estimate of statistical power along the population distribution, apparent patterns identified in any analysis are difficult to interpret with confidence. Consequently, there is a pressing need for the establishment of best practices and new methods for analyzing molecular aging data, discussed below.

### 3.1 Methodology recommendations

There is a great interest in understanding how and why we age, particularly as an increasing number of people are living longer (28). Caring for our aging population requires a deeper understanding of the biological processes that characterize aging and the effects of age on health and wellness. Based on our findings, we offer the following recommendations to improve the integrity of aging research.

#### Critically evaluate the use of LOESS in aging research

LOESS is a powerful tool that can be used to better visualize and understand trends in data; the “bin—summarize—smooth” application of the method has been presented as a useful way to visualize trends in large, complex data (Wickham). However, as with other data visualization techniques, there are important limitations to consider when using LOESS. In particular, LOESS requires dense datasets to produce reliable results. Typically, data are more sparse at the extremes of the predictor distribution; consequently, LOESS curves will be less reliable near the boundaries or in other areas where the population is particularly sparse (20). LOESS curves can also be sensitive to outliers in the data (22). These issues are exacerbated when LOESS curves are used as an input for downstream analyses. Approaches that treat every point along the fitted curve as equally certain may identify apparent changes near the tails of the predictor (age) distribution, even though these patterns are artifacts of overfitting in regions with sparse data. Moreover, LOESS-interpolated values are derived estimates, not independent observations, and should not be analyzed as though they were measured data. Such treatment violates the independence assumptions of most standard statistical tests and can produce spurious statistical significance and misleading biological conclusions. Because LOESS imposes structure on the observed data, statistical tests that are intended to assess the strength of structure in the data will almost always find it in LOESS-interpolated values, resulting in spurious findings.

When used properly, LOESS can be a valuable tool for communicating trends in dense data. To that end, we recommend that LOESS not be used with sparse data where outliers can induce artifactual trends. Further, analysis should not be conducted on LOESS-interpolated data unless the uncertainty in local LOESS estimates—including their induced correlation structure—is propagated throughout the analytical pipeline. For visualizing trends, LOESS plots should also include some measure of uncertainty, such as 95% confidence intervals; methods sections should clearly describe how these measures of uncertainty were calculated. Communicating the level of uncertainty around LOESS estimates will help readers understand how the characteristics of the underlying data may impact the strength of any conclusions drawn.

Although our analyses used LOESS, the artifacts we identified may also arise with other smoothing methods. We were able to detect LOESS-induced artifacts through null simulations, and we recommend that researchers use similar comparisons to assess whether patterns identified in workflows involving smoothing reflect genuine biological signals or artifacts of the analysis.

#### Formulate and conduct hypothesis tests (correctly)

The first step of valid hypothesis testing is the definition of a null hypothesis that envisions what the results would look like absent the phenomenon of interest, and an alternative hypothesis against which it should be tested. Besides providing a clear statistical framework for subsequent claims, the definition of such a hypothesis will eliminate ambiguity around what is meant by “waves of aging.” Currently, there is no consensus on what constitutes an aging “wave.” Formal specifications of null and alternative hypotheses will implicitly define what “waves of aging” are.

The second step of valid hypothesis testing is to compare the observed test statistics to the distribution under the defined null hypothesis, providing an estimate of statistical significance. Test statistics used in aging research tend to be derived from estimators that do not have closed-form null distributions. Permutation testing, which holds the age distribution fixed while randomly shuffling age labels between samples, is a powerful method for empirically constructing a null distribution from the observed data by breaking the dependence between exposure and outcome (in the studies here, age and biomolecule expression levels).

A key assumption of this approach is that data shuffling must be done at the level of exchangeability (*invariance in distribution under finite permutations* (13)) implied by the null hypothesis and study design. In the context of aging research, for observations to be exchangeable under the null hypothesis, permuting the exposure (age) labels must leave the joint distribution of the data unchanged. In other words, the probability distribution of the entire dataset, including the relationships among observations, should be unaffected by shuffling the age labels if there is no true relationship between exposure and outcome (age and expression). Processing data with tools such as LOESS, where each new data point is informed by other data points, induces an index-dependent structure in the data. Therefore, shuffling after such processing will violate the exchangeability assumption and yield an invalid null distribution with misleading permutation results, as occurred in (26).

#### Assess the risk for artifactual findings via simulations and exploratory data analysis

Throughout this paper, we have used arguments from simulated null datasets to investigate the behavior of LOESS, DE-SWAN, and LOESS+DE-SWAN; these simple simulations have demonstrated the key vulnerabilities of these tools. Before using a method, it is important to assess its sensitivity to factors that can induce artifacts in the findings. Often, these sensitivities reveal hidden assumptions that must be satisfied for a result to be valid. Depending on the problem setting, analysts could be able to satisfy these assumptions, but they should always be interrogated. Simulation studies can help reveal possible sources of such artifacts before incorrect conclusions are made from real data. While it is always valuable to have tools that detect true positive signals, it is also important to characterize the circumstances under which they can detect false positive signal. In aging research, statistical inference on trends across the life course is likely to be hampered by data properties such as imbalances in the number of samples available at each age, the overall age distribution, uneven noise, and outliers, among other factors. Basic exploratory data analyses can assist researchers in identifying such factors among their data prior to conducting their main analyses. Perhaps most importantly, simulations in null datasets and exploratory data analysis are both low-overhead tools; the time and cost involved in conducting these steps is small compared to the potential benefits to the robustness and reliability of subsequent analyses.

#### Estimate effects, not significance

Broadly, statisticians recommend using estimated effect sizes with corresponding measures of uncertainty (such as estimated odds ratios with confidence intervals) over measures of significance (*p*-values) to analyze data. Reasons for this are varied, although many of them have been discussed here (30). What becomes challenging about this approach is that it requires the careful propagation of uncertainty in the estimated effect sizes. Doubts about how to best propagate uncertainty are likely what has driven the development of methods based on *p*-values and other related measures of significance. While methods that analyze effect sizes rather than significance are more challenging to develop and implement, such methods will be more robust to many of the structural problems discussed here.

#### Use methods carefully

Methods are rarely as transportable as we wish they were. While methodologists should always design methods to best fit their users’ contexts, users should also be judicious with the methods they operationalize. Analysts should aim to employ methods that have been validated for large sample properties whenever possible and assess whether the necessary assumptions for using the method are satisfied in the context of their research. It is also important to think about a method’s use in its analytical context; using an established method does remove the need for careful study design and confounder analysis.

### 3.2 Concluding remarks

Many published studies have used LOESS and/or DE-SWAN to study molecular or other changes that occur during the aging process (see, for example, Ding et al. (5); Eduthan et al. (6); Lai et al. (15); Lehallier et al. (17); Liu et al. (18, 19); Shen et al. (26, 27); Yin et al. (31); You et al. (32)). The analysis presented here has not considered each of these studies in detail and, therefore, cannot be taken as a direct refutation of their findings. Our results, however, do cast doubt on their conclusions. Further, although nothing in our analysis disproves the existence of nonlinearities in the aging process, we have shown that patterns similar to those reported can emerge from random noise. We believe this plainly demonstrates that some of the most commonly used tools in molecular aging research cannot reliably identify molecular shifts throughout the aging process. Recent work by Che et al. independently arrived at similar conclusions (1). Developing new ways to overcome the methodological limitations we have identified and establishing best practices will be crucial for characterizing the molecular factors that contribute to the biological process of aging.

In analyzing cross-sectional data, we identified non-uniform age distribution as one possible source of artifactual waves of aging, and showed that the risk of certain artifacts, such as non-linear trajectories determined by LOESS, may be reduced in the case of a uniform age distribution (Figure 2B). However, a uniform age distribution does not eliminate the possibility that such artifacts can arise. Artifactual waves of aging data can arise from heteroskedasticity even when the age distribution is uniform (Figure 10), and outliers can also induce apparent waves when the age distribution is uniform (Supplementary Figure **??**). The age distribution used in a study must be given careful consideration in light of the questions one is seeking to answer; a uniform distribution of ages across the lifespan may exacerbate other confounding factors, such as survivor bias, that must be appropriately addressed. Longitudinal data are preferable to cross-sectional data for several reasons. However, most of the methodological problems discussed here would also be problematic in a longitudinal setting. Major studies such as NHANES and the Baltimore Longitudinal Study of Aging have biobanking initiatives that are collecting such data, but the number of time points assessed for each individual in the study is relatively small when compared to large cross-sectional cohorts. This proposes methodological challenges as it can become difficult to accurately model aging as a continuum when the sampling points are so discrete. Therefore, in the case of both uniformly sampled cross-sectional data and longitudinal analyses, the best approach forward will be new methodological development based on the principles outlined in section 3.1.

We have introduced a range of methodological recommendations and areas for further investigation above, but our central point is this: while the field of aging science has dedicated substantial resources to achieving excellence in the establishment of meticulously curated biorepositories for studying molecular characteristics of aging, investment in methodological development lags far behind. This is an exciting moment for aging science: the vast amount of available molecular aging data is ready for the next generation of aging methodologists to decipher it with new and powerful tools. We have written this paper not as a critique, but as a call to action towards discovery of real, meaningful scientific insights that will improve the universal human experience of aging.

## 4 Online Methods

### 4.1 Methods: Artifact 1

#### 4.1.1 Simulation example on a single gene

We simulated a population of 100 individuals with normally distributed ages (mean = 55, standard deviation = 7). We then simulated expression of a single molecule for these individuals by drawing a sample of size 100 from a standard normal distribution, independent of age. A span of 0.75 was used for LOESS and a sliding window width of 20 years was used for DE-SWAN.

### 4.2 Methods: Artifact 2

#### 4.2.1 Waves of aging appear in null data and depend on the age distribution

For each age distribution, we simulated expression levels for *p* = 1000 genes by drawing independent samples from a standard normal distribution. We then ran LOESS+DE-SWAN with a window size of 20 years and an increment of one year, following the analysis of Lehallier et al. (17) and Shen et al. (26). The initial window center was set at the minimum age plus 10 years and the LOESS span was set to 0.6. As in Shen et al. (26), we generated LOESS-interpolated data by sampling the LOESS curve along a grid of points 0.5 years apart across the age distribution. We ran the modified DE-SWAN algorithm on these LOESS-interpolated data to compute the number of molecules detected as significantly changing (FDR *<* 0.05) at each time point. To assess variability, we repeated this process 100 times per age distribution.

For the iPOP replication analysis, we paired the iPOP age distribution with 8,556 simulated genes, each following a standard normal distribution independent of age. Following Shen et al. (26), we adjusted the first window to be centered at age 40 and to include data ranging from age 25 to 50. The remainder of the windows were centered along the sequence (41, 42*,…,* 65) with width 20, considering data points located within *±* 10 years from each window center. To confirm that a fixed choice of LOESS span was not unduly influencing the observation of artifactual waves of aging, we applied the LOESS span optimization strategy adopted by Shen et al. (26) (based on the function found at https://github.com/jaspershen-lab/ipop_aging/blob/main/1-code/100-tools.R#L203). We repeated this process 50 times.

### 4.3 Methods: Artifact 3

#### 4.3.1 Sample Age Distribution

Data were simulated for *n* = 1000 people for *p* = 2000 molecules. Simulations were run 100 times. The first sliding window was centered at 25 and the last sliding window was centered at 75; window lengths are described below. All data were simulated from a linear model. For each observation *i* = 1*,…, n* and *j* = 1*,…, p*

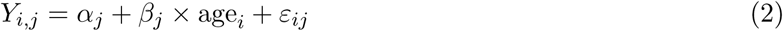

where 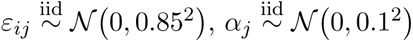 and 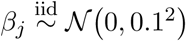.

For the uniform age distribution, 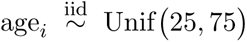. For the normal age distribution, age 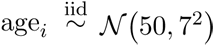. Both the uniform and normal distribution simulations were run with an interval length of 10 years (20-years of data were used for each sliding window). For the bimodal (20-yr) age distribution, age 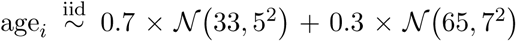. For the bimodal (10-yr) age distribution, 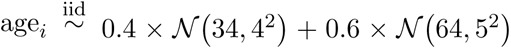. The bimodal distribution simulations were each run with a window length of 20 and 10 years, respectively.

#### 4.3.2 Heteroskedasticity

Data were simulated similarly to the Sample Age Distribution simulations (Online Methods 4.3.1). However, the ages of each simulation were drawn from 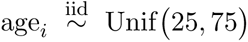 and this procedure did not change from simulation to simulation. Furthermore, the *ε_ij_* term is generated differently in equation 2. For each age, the standard deviation of the random component *ε_ij_* is multiplied by a scalar. For the “High Noise in Center” simulations, the multiplier was chosen so that the multiplier increases linearly until the age midpoint and then decreases linearly by the same rate (see Figure 10). For the “High Noise at Ends” simulations, the sign of the multiplier is flipped. In the “Noise Increase” simulations, the multiplier was simulated to increase exponentially, and for the “Noise Decrease” simulations, the multiplier was simulated to decrease exponentially. Parameter values can be found in https://github.com/QuackenbushLab/artifactual-waves-of-aging.

### 4.4 Methods: Literature Search

To gain an understanding of the breadth of this issue in the literature, we conducted complementary literature searches in Google Scholar, Scopus, and ScienceDirect and integrated the results.

#### 4.4.1 Google Scholar

To collect Google Scholar search results, we used the freely available Publish or Perish software (11), version 8.19.5300 on MacOS. We conducted a Google Scholar search limited to the years 2019-2026 using the search term:

(loess OR de-swan OR deswan) AND aging lehallier.

We conducted this search on September 8, 2026; the search yielded 168 results.

#### 4.4.2 Scopus

Unlike Google Scholar, Scopus does not provide a full-text search, but rather queries attributes such as abstracts, references, and metadata. For this reason, we broadened our Google Scholar search term to include the words “nonlinear” or “non-linear”, which we felt were more likely to appear in titles, abstracts, and keywords than LOESS or DE-SWAN. Additionally, we removed “lehallier” from the search term and instead required that Lehallier et al. 2019 be cited. The final search query was:

TITLE-ABS-KEY (”LOESS” OR “de-swan” OR “deswan” OR “nonlinear” OR “non-linear”) AND REFTITLE (”Undulating changes in human plasma proteome profiles across the lifespan”)

We conducted our Scopus search on September 9, 2026; the search yielded 44 results.

#### 4.4.3 ScienceDirect

Like Google Scholar, ScienceDirect provides a full-text search and allows the specification of references. We therefore used nearly the same search term as our Google Scholar search, but omitted “lehallier” and instead required that Lehallier et al. 2019 be cited. The final ScienceDirect query was:

Find articles with these terms: (“loess” OR “de-swan” OR “deswan”) AND aging References: Undulating changes in human plasma proteome profiles across the lifespan

We conducted our ScienceDirect search on September 9, 2026; the search yielded 23 results.

#### 4.4.4 Cleaning and integrating Google Scholar, Scopus, and ScienceDirect searches

We first filtered for any “duplicate” publications that were represented by more than one search result (for example, a preprint and a published manuscript were both recovered). There were four duplicates in the Google Scholar results and no duplicates in either the Scopus or ScienceDirect searches. In the case of duplicates, the record with the best (minimum) Google Scholar rank was retained and the others were discarded.

There was no consistently reported document identifier (DOI, PMID, PMCID) across all three databases. To join results, we relied on title similarity. Similarity was calculated using the stringdist R package with the Levenshtein distance (method = “lv”), and up to five mismatches were allowed. We first performed a fuzzy full join of the Google Scholar and Scopus results, then performed two fuzzy joins of the Scholar-Scopus result with the ScienceDirect result: a fuzzy full join based on Scholar-ScienceDirect title similarity, and a fuzzy inner join based on Scopus-ScienceDirect title similarity to capture any records detected with Scopus and ScienceDirect that were not in the Google Scholar results. Concatenating these two data frames yielded 189 records, the majority of which were found only by Google Scholar (Supplementary Table **??**).

#### 4.4.5 Methods for screening search results

We screened each of the 189 articles found by the union of the three searches to better understand the accuracy of these queries in identifying potentially problematic publications.

Two humans (MC and KHS) screened each article to determine (1) if Lehallier et al. (17) was cited, (2) if LOESS and/or DE-SWAN was used in the methodology, and (3) if the main topic is related to aging or age-related diseases. As an exploratory exercise, two agent-based screens were also conducted, one by a GPT-5–family Codex agent running in the Codex desktop app, and one by Claude Cowork in the Claude desktop app, configured for Claude Sonnet 5 (knowledge cutoff of January 2026; the agent used web search to fill in anything more recent).

The following prompt was used:

For each record, determine the following three things. 1. Does the record cite the Nature Medicine paper “Undulating changes in human plasma proteome profiles across the lifespan” by Lehallier et al. 2019? Answer yes or no. 2. Is LOESS and/or DE-SWAN used in the methodology? Answer yes, no, or uncertain. 3. Is the main topic of the paper related to aging or age-related diseases? Answer yes or no. Add any additional notes that would help me understand your reasoning. For records linked to papers that are not written in English or that are duplicated elsewhere in the list, please write OMIT (except for the first instance in duplicated papers). Output your results in the appropriate columns of the uploaded spreadsheet and provide the results back as a spreadsheet. Read the papers carefully and check online supplements if necessary to resolve any ambiguity.

## 5 Author Contributions

MC and KHS conceptualized the study and conducted the experiments. MC, KHS, and JQ drafted the manuscript. XS and MPS clarified the original analytical workflow, independently reviewed the reanalysis, contributed to the interpretation of affected conclusions, and revised the manuscript for methodological and biological accuracy. JWM and JQ provided supervision. All authors reviewed and edited the manuscript.

## 6 Funding and Disclosures

MC was supported by the National Institute of General Medical Sciences of the National Institutes of Health under award number T32GM135117.

KHS was supported by the National Heart, Lung, and Blood Institute of the National Institutes of Health under award number K25HL175222.

JWM has no funding or disclosures to declare in relation to this work.

MPS is a cofounder and scientific advisor of Crosshair Therapeutics, Exposomics, Filtricine, Fodsel, iollo, InVu Health, January AI, Marble Therapeutics, Mirvie, Next Thought AI, Orange Street Ventures, Personalis, Protos Biologics, Qbio, RTHM, SensOmics. MPS is a scientific advisor of Abbratech, Applied Cognition, Enovone, Jupiter Therapeutics, M3 Helium, Mitrix,Neuvivo, Onza, Sigil Biosciences, TranscribeGlass, WndrHLTH, Yuvan Research. MPS is a cofounder of NiMo Therapeutics. MPS is an investor and scientific advisor of R42 and Swaza.MPS is an investor in Repair Biotechnologies.

XS has no funding or disclosures to declare in relation to this work.

JQ was supported by the National Cancer Institute of the National Institutes of Health under award number R35CA220523.

The content is solely the responsibility of the authors and does not necessarily represent the official views of the National Institutes of Health.

## Notes

https://github.com/QuackenbushLab/artifactual-waves-of-aging

### Competing Interest Statement

Michael Snyder is a cofounder and scientific advisor of Crosshair Therapeutics, Exposomics, Filtricine, Fodsel, iollo, InVu Health, January AI, Marble Therapeutics, Mirvie, Next Thought AI, Orange Street Ventures,
Personalis, Protos Biologics, Qbio, RTHM, SensOmics. He is a scientific advisor of Abbratech, Applied Cognition, Enovone, Jupiter Therapeutics, M3 Helium, Mitrix, Neuvivo, Onza, Sigil Biosciences, TranscribeGlass, WndrHLTH, Yuvan Research. MPS is a cofounder of NiMo Therapeutics. He is also an investor and scientific advisor of R42 and Swaza.MPS is an investor in Repair Biotechnologies.

### Summary of Updates

We have updated the manuscript to address referee concerns and in response to a query from the editor of the journal to which the manuscript was submitted. The changes are minor except for an analysis of the published literature to address a question regarding the possible extent ot the problem caused by the use of DE-SWAN to analyze molecular aging data.

## References

[1] Che, A., Shah, P., Tanner, K. T., and Cohen, A. A. (2026). De-swan analysis of biomarker trajectories over the life course is highly sensitive to age distribution and model parameterization. bioRxiv, pages 2026–09.

[2] Chen, R., Mias, G. I., Li-Pook-Than, J., Jiang, L., Lam, H. Y., Chen, R., Miriami, E., Karczewski, K. J., Hariharan, M., Dewey, F. E., et al. (2012). Personal omics profiling reveals dynamic molecular and medical phenotypes. Cell, 148(6):1293–1307.

[3] Cleveland, W. S. and Devlin, S. J. (1988). Locally weighted regression: an approach to regression analysis by local fitting. Journal of the American Statistical Association, 83(403):596–610.

[4] Cummings, S. R., Newman, A. B., Coen, P. M., Hepple, R. T., Collins, R., Kennedy, MS, K., Danielson, M., Peters, K., Blackwell, T., Johnson, E., Mau, T., Shankland, E. G., Lui, L.-Y., Patel, S., Young, D., Glynn, N. W., Strotmeyer, E. S., Esser, K. A., Marcinek, D. J., Goodpaster, B. H., Kritchevsky, S., and Cawthon, P. M. (2023). The study of muscle, mobility and aging (somma): A unique cohort study about the cellular biology of aging and age-related loss of mobility. The Journals of Gerontology: Series A, 78(11):2083–2093.

[5] Ding, Y., Zuo, Y., Zhang, B., Fan, Y., Xu, G., Cheng, Z., Ma, S., Fang, S., Tian, A., Gao, D., Xu, X., Wang, Q., Jing, Y., Jiang, M., Xiong, M., Li, J., Han, Z., Sun, S., Wang, S., He, F., Yang, J., Qu, J., Zhang, W., and Liu, G.-H. (2025). Comprehensive human proteome profiles across a 50-year lifespan reveal aging trajectories and signatures. Cell, 188(20):5763–5784.e26.

[6] Eduthan, N. P., Donovan, M. G., Araya, P., Rachubinski, A. L., Sullivan, K. D., Galbraith, M. D., and Espinosa, J. M. (2025). An integrated multi-omic natural history study of human development, sexual dimorphism, and the effects of trisomy 21. Nature Communications, 16(1):8346.

[7] Farrar, T., Blignaut, R., Luus, R., and Steel, S. (2025). A review and comparison of methods of parameter estimation and inference for heteroskedastic linear regression models. Journal of Applied Statistics, 52(16):3091–3120. eprint: 10.1080/02664763.2025.2496719.

[8] Ferrucci, L. (2008). The Baltimore Longitudinal Study of Aging (BLSA): A 50-Year-Long Journey and Plans for the Future. The journals of gerontology. Series A, Biological sciences and medical sciences, 63(12):1416–1419.

[9] Good, P. I. (2002). Extensions of the concept of exchangeability and their applications. Journal of Modern Applied Statistical Methods, 1(2):34.

[10] Griffiths, W. E. (2003). Heteroskedasticity. In A Companion to Theoretical Econometrics, pages 82–100. John Wiley & Sons, Ltd. Section: 4 eprint: https://onlinelibrary.wiley.com/doi/pdf/10.1002/9780470996249.ch5.

[11] Harzing, A. (2007). Publish or Perish.

[12] Jacoby, W. G. (2000). Loess: a nonparametric, graphical tool for depicting relationships between variables. Electoral Studies, 19(4):577–613.

[13] Kallenberg, O. (2021). Foundations of Modern Probability. Springer. Chapter 25, pages 557–585.

[14] Kedlian, V. R., Donertas, H. M., and Thornton, J. M. (2019). The widespread increase in interindividual variability of gene expression in the human brain with age. Aging, 11(8):2253–2280.

[15] Lai, W., Feng, Q., Lei, W., Xiao, C., Wang, J., Zhu, Y., Mao, L., Zhu, Y., He, J., Li, Y., et al. (2025). Deciphering immunosenescence from child to frailty: transcriptional changes, inflammation dynamics, and adaptive immune alterations. Aging Cell, 24(7):e70082.

[16] Lehallier, B. (2026). DEswan: Differential Expression Sliding Window ANalaysis. R package version 0.0.0.9001, commit be2dd3c64614cd288a1afabc89bd8af40bbfced2.

[17] Lehallier, B., Gate, D., Schaum, N., Nanasi, T., Lee, S. E., Yousef, H., Moran Losada, P., Berdnik, D., Keller, A., Verghese, J., Sathyan, S., Franceschi, C., Milman, S., Barzilai, N., and Wyss-Coray, T. (2019). Undulating changes in human plasma proteome profiles across the lifespan. Nature Medicine, 25(12):1843–1850.

[18] Liu, W.-S., You, J., Chen, S.-D., Zhang, Y., Feng, J.-F., Xu, Y.-M., Yu, J.-T., and Cheng, W. (2025a). Plasma proteomics identify biomarkers and undulating changes of brain aging. Nature Aging, 5(1):99–112.

[19] Liu, X., Liang, T., Zhao, R., Zhu, M., Huang, B., Huang, X., and Ni, F. (2025b). A Global Metabolomic and Lipidomic Landscape of Human Plasma Across the Lifespan.

[20] Loader, C. and Loader, C. (1999). Local Regression and Likelihood. Statistics and Computing. Springer New York.

[21] Nichols, E. and Hayes-Larson, E. (2025). Common methodological issues in observational epidemiological studies of older adults. BMJ Medicine, 4(1):e001332.

[22] NIST/SEMATECH (2023). LOESS (aka LOWESS). NIST/SEMATECH e-Handbook of Statistical Methods. Section 4.1.4.4, Process Modeling.

[23] Piehl, N., Olst, L. v., Ramakrishnan, A., Teregulova, V., Simonton, B., Zhang, Z., Tapp, E., Channappa, D., Oh, H., Losada, P. M., Rutledge, J., Trelle, A. N., Mormino, E. C., Elahi, F., Galasko, D. R., Henderson, V. W., Wagner, A. D., Wyss-Coray, T., and Gate, D. (2022). Cerebrospinal fluid immune dysregulation during healthy brain aging and cognitive impairment. Cell, 185(26):5028–5039.e13.

[24] Poulton, R., Moffitt, T. E., and Silva, P. A. (2015). The Dunedin Multidisciplinary Health and Development Study: overview of the first 40 years, with an eye to the future. Social Psychiatry and Psychiatric Epidemiology, 50(5):679–693.

[25] Pugh, S. L. and Molinaro, A. (2016). The nuts and bolts of hypothesis testing. Neuro-Oncology Practice, 3(3):139–144.

[26] Shen, X., Wang, C., Zhou, X., Zhou, W., Hornburg, D., Wu, S., and Snyder, M. P. (2024). Nonlinear dynamics of multi-omics profiles during human aging. Nature Aging, 4(11):1619–1634.

[27] Shen, X., Wu, B., Jiang, W., Li, Y., Zhang, Y., Zhao, K., Nie, N., Gong, L., Liu, Y., Zou, X., Liu, J., Jin, J., and Ouyang, H. (2022). Scale bar of aging trajectories for screening personal rejuvenation treatments. Computational and Structural Biotechnology Journal, 20:5750–5760.

[28] Shreya, D., Fish, P. N., and Du, D. (2025). Navigating the future of elderly healthcare: a comprehensive analysis of aging populations and mortality trends using National Inpatient Sample (NIS) data (2010-2024). Cureus, 17(3):e80442.

[Wickham] Wickham, H. Bin-summarise-smooth: a framework for visualising large data. Technical Report. 2013.

[30] Williams, S., Carson, R., and Tóth, K. (2023). Moving beyond P values in The Journal of Physiology: A primer on the value of effect sizes and confidence intervals. The Journal of Physiology, 601(23):5131–5133. eprint: https://physoc.onlinelibrary.wiley.com/doi/pdf/10.1113/JP285575.

[31] Yin, J., Ibrahim, S., Petersen, F., and Yu, X. (2021). Autoimmunomic Signatures of Aging and Age-Related Neurodegenerative Diseases Are Associated With Brain Function and Ribosomal Proteins. Frontiers in Aging Neuroscience, 13:679688.

[32] You, J., Cui, X.-H., Chen, Y.-L., Wang, Y.-X., Li, H.-Y., Qiang, Y.-X., Cheng, J.-Y., Deng, Y.-T., Guo, Y., Ren, P., et al. (2025). Mapping the plasma metabolome to human health and disease in 274,241 adults. Nature Metabolism, 7(11):2366–2384.

