## Supplemental Materials for "LOESS and DE-SWAN can induce artifactual “waves” of molecular aging"

**S3 Authors' Note** **5**

| Description | Scholar | Scopus | SciDir | N |
|---|---|---|---|---|
| Scholar Only | X | O | O | 130 |
| Scopus Only | O | X | O | 19 |
| SciDir Only | O | O | X | 4 |
| Scholar and Scopus | X | X | O | 17 |
| Scholar and SciDir | X | O | X | 11 |
| Scopus and SciDir | O | X | X | 2 |
| Scholar and Scopus and SciDir | X | X | X | 6 |
| Total | | | | 189 |

Supplementary Table S1: Breakdown of the sources of the 189 total records obtained by querying Google Scholar via Publish or Perish ("Scholar"), Scopus, and Science Direct ("SciDir").

| Description | N |
|---|---|
| Scholar Only | 37 |
| Scopus Only | 2 |
| SciDir Only | 0 |
| Scholar and Scopus | 11 |
| Scholar and SciDir | 7 |
| Scopus and SciDir | 2 |
| Scholar and Scopus and SciDir | 4 |
| Total | 63 |

Supplementary Table S2: Sources of the 63 papers confirmed to meet all three screening criteria by both MC and KHS.

### S1 Literature Search

#### S1.1 Literature Search Characteristics

#### S1.2 Results of literature query

##### S1.2.1 Consensus by human reviewers

While both human reviewers agreed in 100 percent of records on whether or not Lehallier et al. (2019) was referenced by each record, the AI reviewers did not do well with this task, notably missing clear citations (Supplementary Figure S2A). It was often challenging to answer the second question, "Is LOESS and/or DE-SWAN used in the methodology?", even after manual review of the manuscripts. All four reviewers (MC, KHS, Codex, Claude) had a substantial number of "uncertain" labels; MC: 39/189, KHS: 47/189, Codex: 18/189, Claude: 23/189. Moreover, the agreement as to which papers were uncertain was sparse (Supplementary Figure S2B). Generally, all four reviewers agreed on whether the paper's primary topic was aging or age-related disease (Supplementary Figure S2C).

Based on the poor performance of the AI searches on what we perceived to be an objective question (citation) and the overall inconsistency of their results throughout, we did not use the AI results to inform our final conclusions on this matter. Rather, we sought consensus by the human reviewers.

Of the original 189 records, 63 unique manuscripts were confirmed to be of primary concern based on meeting all three criteria by as assessed by both human reviewers (MC and KHS). The breakdown of these records by source is shown in Supplementary Table S2. The number of such manuscripts per year has been rapidly increasing since 2024 (Supplementary Figure S1), emphasizing the timeliness of this issue and the critical importance of vigilance among editors and peer reviewers in screening for these methodological concerns.

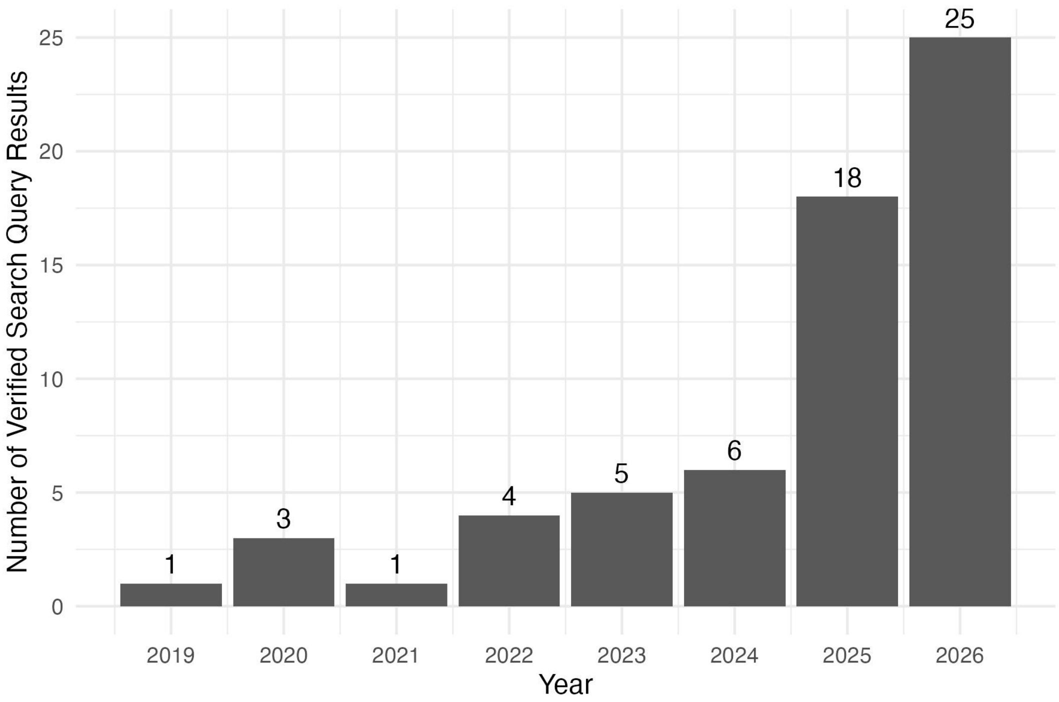

Supplementary Figure S1: Number of published papers/preprints identified as high concern based on the literature search, verified by two independent screeners (KHS and MC).

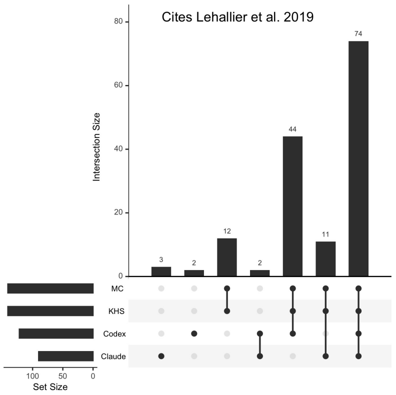

(A) Determination of "Yes" for citation of Lehallier et al. (2019).

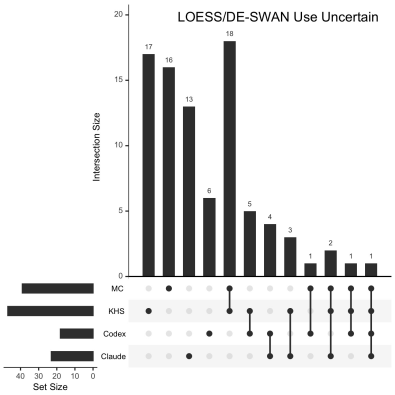

(B) Determination of "uncertain" for the use of LOESS and/or DE-SWAN.

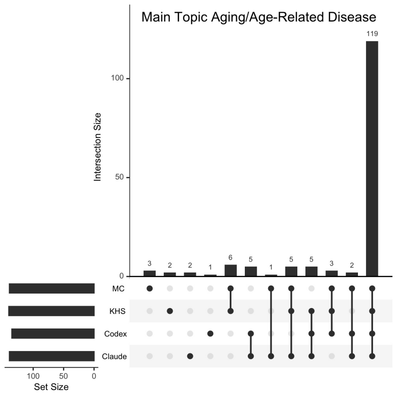

(C) Determination of "Yes" for aging or aging-related disease as topic.

Supplementary Figure S2: Summary characteristics of consensus between two human reviewers (MC, KHS) and two AI reviewers (Codex, Claude).

### S2 Additional notes regarding simulations

#### S2.1 Choice of cluster count in "Smooth-Cluster-Interpret" workflow

As the cut point of the dendrogram is adjusted to yield greater numbers of clusters, the specificity of the cluster shapes can increase, leading to erroneous conclusions about hallmark molecular trajectories of aging. Figure S3 illustrates this issue. For $K = 2$ clusters, the null trajectories are generally appropriately grouped into a cluster whose centroid exhibits a null association with aging. As $K$ increases, centroids with peaks at the younger or older end of the age continuum are observed. Because these data have no true association with age, such characteristic trajectories are fitting noise rather than signal.

`figs/random_noise_sims/clustering/alter_cluster_counts.pdf`

Supplementary Figure S3: Examples of the effect of different choices of $K$ on characteristic trajectories detected in random data for the Gaussian distribution described in Online Methods ??.

##### S2.2 Injected Outliers in DE-SWAN

Worth a brief discussion is the impact that outliers can have on DE-SWAN curves. Standard ordinary least squares fitted linear regression is not robust to outliers. Therefore, the findings of standard regression-based tests used in DE-SWAN will also be susceptible to outliers. For DE-SWAN, the presence of outliers can be particularly problematic in small datasets where outliers are unevenly distributed throughout the age distribution. Outliers modulate statistical power throughout the age distribution through either artificially increasing or decreasing the effect size. In small datasets, several of these outlier observations can have a significant impact on the observed signal. We conducted simulation studies to assess this issue as follows. Results are shown in Figure S4.

Data were simulated for $n = 200$ people for $p = 1500$ molecules. Simulations were run 50 times. The first sliding window was centered at 25 and the last sliding window was centered at 75. Data were again simulated similarly to the Sample Age Distribution simulations (Supplementary Section **??**). The ages of each simulation were also drawn from $\text{age}_i \overset{\text{iid}}{\sim} \text{Unif}(25, 75)$ and this procedure did not change from simulation to simulation. In Figure S4, 5 outliers were simulated to be located in the center of the age distribution. Outliers were generated from the model (**??**) with $\varepsilon_{ij} \overset{\text{iid}}{\sim} \mathcal{N}(0, (5 \times 0.85)^2)$ to increase their variability.

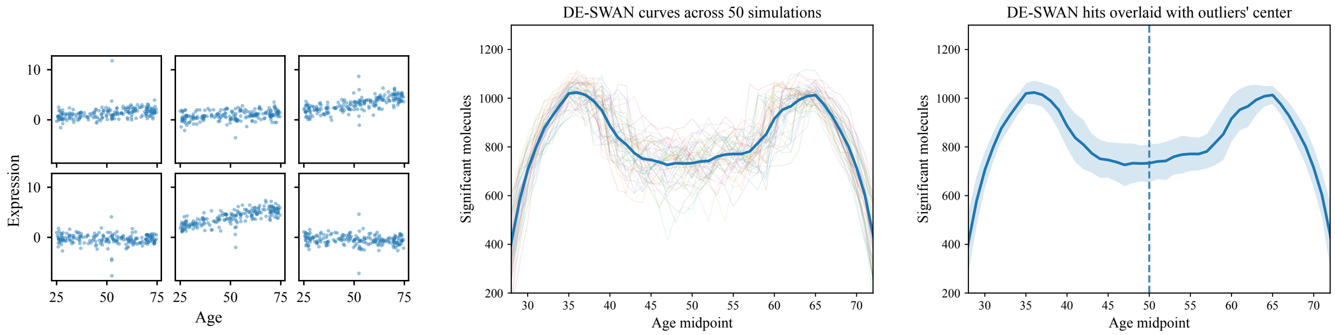

Supplementary Figure S4: Effects of outliers on DE-SWAN results. Each dataset was simulated for 200 observations and 1,500 molecules. Results from 50 simulations are shown. The first panel shows six randomly drawn molecular expression profiles with five observations designated as outliers. Between simulation settings, outliers were chosen to be at the center of the age distribution. The second column shows DE-SWAN curves plotted for 50 simulations, with the mean curve plotted in blue. The third column shows the mean DE-SWAN curve plotted with a $\pm 1$ SD band (solid blue). The dashed blue line shows the where outliers were positioned. Simulation details can be found in S2.2.

#### S3 Authors' Note

In this article, we provide a critical evaluation of how LOESS and DE-SWAN are used to analyze data in aging research. We specifically comment on how the LOESS+DE-SWAN pipeline employed by Shen et al. (2024) is invalid for the analysis of these data. We caution against using these methods to analyze age-related molecular profiles and highlight the need for appropriate methods in this important area of research.

We would be remiss not to comment on the importance of reproducible research and on the positive example set by Shen et al. (2024). Open-source software and truly available data are cornerstones of modern science as they make our work transparent, testable, and ultimately trustworthy. Reproducibility and falsifiability are not abstract philosophical ideals—they are operational requirements for progress in any quantitative discipline. When we share code, workflows, and data, we enable others to evaluate our assumptions, replicate our findings, and extend our results in ways that would otherwise be impossible. Shen et al. (2024) are to be commended for their unflinching adoption of open-source principles.

Just as importantly, the identification of errors in published work should never be viewed as a weakness of the scientific enterprise. It is precisely the opposite. Science advances because results are challenged, reanalyzed, and sometimes corrected in the open. That process—rigorous scrutiny followed by refinement—is the mechanism by which we move closer to the truth. In that sense, openness is not simply a cultural preference; it is the infrastructure that makes cumulative scientific progress possible.

Many published studies in aging employ LOESS and DE-SWAN, either separately or together. We initially planned to conduct a more comprehensive analysis across many published studies but, unfortunately, we found that obtaining data and code frequently imposed difficulties that would have significantly delayed the completion of this manuscript. The fact that we could clearly delineate these issues is due to the adherence of Shen et al. (2024) to the principles of open science. This should not be taken as a sign of poor science or a failure of peer review, but a testament to the value of open science principles.

Furthermore, it is worth noting that finding fault with an analysis in a published paper need not be an adversarial process. When we discovered that the analytical pipeline used by Shen et al. (2024) had detected artifactual trends in the data, we conducted an exhaustive analysis of the method to understand the source of the problem. We contacted the senior and first authors of the paper and arranged a meeting with them to share our analyses. We concluded by providing them with a detailed summary of our results, allowing them time to investigate the issue themselves. We independently met with the editor of the journal in which the study was published to clarify that this was not a "disagreement between authors about interpretation" but rather the identification of an honest error in the analyses. This manuscript is due, in part, to encouragement from the editor on the importance of raising awareness of the issue. Scientists are humans who make mistakes—and good scientists acknowledge those errors when pointed out, as have Shen et al. (2024).

Finally, it should be acknowledged that the trends reported by Shen et al. (2024) are biologically plausible given that we know aging has profound effects on human health and vitality, that many of those changes correlate with menopause/andropause, and that changes ranging from gray hair and wrinkled skin to altered metabolism and immunity often occur in the 40s and 60s. Thus, it is possible that the study's scientific conclusions will be borne out, even though the current data and methods are insufficient to demonstrate them.

### References

Lehallier, B., Gate, D., Schaum, N., Nanasi, T., Lee, S. E., Yousef, H., Moran Losada, P., Berdnik, D., Keller, A., Verghese, J., Sathyan, S., Franceschi, C., Milman, S., Barzilai, N., and Wyss-Coray, T. (2019). Undulating changes in human plasma proteome profiles across the lifespan. *Nature Medicine*, 25(12):1843–1850.

Shen, X., Wang, C., Zhou, X., Zhou, W., Hornburg, D., Wu, S., and Snyder, M. P. (2024). Nonlinear dynamics of multi-omics profiles during human aging. *Nature Aging*, 4(11):1619–1634.
